# Transcriptional regulators ensuring specific gene expression and decision making at high TGFβ doses

**DOI:** 10.1101/2024.04.23.590740

**Authors:** Laura Hartmann, Panajot Kristofori, Congxin Li, Kolja Becker, Lorenz Hexemer, Stefan Bohn, Sonja Lenhardt, Sylvia Weiss, Björn Voss, Alexander Loewer, Stefan Legewie

## Abstract

TGFβ-signaling regulates cancer progression by controlling cell division, migration and death. These outcomes are mediated by gene expression changes, but the mechanisms of decision making towards specific fates remain unclear. Here, we combine SMAD transcription factor imaging, genome-wide RNA sequencing and morphological assays to quantitatively link signaling, gene expression and fate decisions in mammary epithelial cells. Fitting genome-wide kinetic models to our time-resolved data, we find that the majority of TGFβ target genes can be explained as direct targets of SMAD transcription factors, whereas the remainder show signs of complex regulation, involving delayed regulation and strong amplification at high TGFβ doses. Knockdown experiments followed by global RNA sequencing revealed transcription factors interacting with SMADs in feedforward loops to control delayed and dose-discriminating target genes, thereby reinforcing the specific epithelial-to-mesenchymal transition at high TGFβ doses. We identified early repressors, preventing premature activation, and a late activator, boosting gene expression responses for a sufficiently strong TGFβ stimulus. Taken together, we present a global view of TGFβ-dependent gene regulation and describe specificity mechanisms reinforcing cellular decision making.

## Introduction

The TGFβ signaling pathway controls embryonic development and adult tissue homeostasis. Its dysregulation has been linked to various diseases such as fibrosis and cancer (Akhurst and Hata, 2012). During cancer progression TGFβ plays a dual role. In early stages of tumorigenesis, the TGFβ signaling pathway has tumor suppressive properties by promoting apoptosis and inhibiting cell proliferation. In later stages, it acts as a tumor promoter by preventing cell cycle arrest and initiating epithelial-mesenchymal-transition (EMT), which is a hallmark of cancer, to acquire migratory properties by down regulation of epithelial and upregulation of mesenchymal markers (Ikushima and Miyazono, 2010), (Siegel and Massagué, 2003), (Xu, Lamouille and Derynck, 2009). The underlying molecular mechanisms controlling the shift from tumor suppressor to tumor promoter function are currently unclear but may involve differential expression of transcriptional factors (TFs) and alterations in the temporal dynamics of signaling (Mullen *et al*., 2011), (Piek *et al*., 2001).

TGFβ functions as an extracellular cytokine that triggers intracellular signaling by binding to and activating its transmembrane serine/threonine kinase receptors, TGFβRI and TGFβRII. The activated receptors phosphorylate cytoplasmic signal transducers, SMAD2, and SMAD3, which subsequently heterotrimerize with SMAD4, translocate to the nucleus, and function as TFs by binding to the promoters of target genes (Feng and Derynck, 2005). The SMAD2/3/4 complex binding has low affinity to SMAD-binding-elements (SBE) in promoter regions and transcriptional regulation by SMAD is therefore dependent on additional proteins such as TFs and epigenetic regulators (Morikawa *et al*., 2013), (Shi *et al*., 1998), (Hill, 2016). SMADs utilize their Mad Homology 1 (MH1) and Mad Homology 2 (MH2) domains to bind to DNA and interact with co-repressors and co-activators, respectively (Feng and Derynck, 2005), (Shi *et al*., 1998). Known co-regulators that guide the SMAD-complexes to distinct promoter sequences and regulate target gene expression, include TFs such as ATF3, RUNX1, TGIF, SKIL, or SKI, and epigenetic regulators like HDAC1/3/4/5/6 and GCN5 (Miyazawa *et al*., 2002), (Ross and Hill, 2008). Upon TGFβ stimulation, SMADs can induce the expression of their collaborating TFs. JUNB, RUNX1, SNAI1, and SNAI2 thereby establishing feed-forward-loops (FFL), in which SMAD controls target genes both directly via promoter binding and indirectly via co-factor up-regulation (Sundqvist *et al*., 2013), (Peinado, Quintanilla and Cano, 2003), (Sundqvist *et al*., 2018).

Current evidence suggests that the temporal dynamics of SMAD signaling play an important role in cellular decision making. Using time-lapse microscopy of individual MCF10A cells, we found that transient SMAD2 dynamics are associated with increased cell motility, while sustained SMAD2 dynamics additionally induce cell cycle arrest and further reinforced cellular motility towards EMT (Strasen *et al*., 2018; Bohn *et al*., 2023). Comparable results were observed in pancreatic cancer cell lines, where a sustained SMAD signal was required for cell cycle arrest (Nicolás and Hill, 2003). Additionally, in myoblasts TGFβ pulses as opposed to continuous stimulation, induced target gene expression and fate regulation (Sorre *et al*., 2014). Studies on other signaling pathways have shown that the strength and duration of intracellular signals generally play a pivotal role in influencing the decisions cells make regarding their fate, e.g., in MAPK-induced proliferation vs. neuronal differentiation of PC12 cells, or during p53-induced DNA repair vs. cellular senescence (Behar and Hoffmann, 2010; Purvis and Lahav, 2013), (Marshall, 1995), (Purvis *et al*., 2012). To induce specific cell fates, different gene expression programs need to be induced depending on the temporal dynamics of signaling. Prior studies combined RNA sequencing with mathematical modelling to investigate EMT regulators upon TGFβ-signaling (Deshmukh *et al*., 2021). However, it has been a matter of active research how gene expression networks distinguish input signals e.g., through epigenetic regulation mechanisms as well as feedback- and feedforward regulation by TFs resulting in different phenotypic outcomes (Mangan and Alon, 2003), (Weidemüller *et al*., 2021). For instance, systems biological approaches combining quantitative experiments with dynamic modelling have shown that coherent FFL, where both inputs directly and indirectly induce gene expression networks, allow for decoding the amplitude and duration of input signals (Alon, 2007). Furthermore, signal decoding at the level of target gene expression may be related to mRNA half-life (Uhlitz *et al*., 2017).

For TGFβ-SMAD signaling, systems biological studies investigated the molecular mechanisms shaping temporal dynamics of signal transmission from the cell membrane to the nucleus, or focused on the downstream target gene expression level (Zi *et al*., 2011), (Zi, Chapnick and Liu, 2012), (Strasen *et al*., 2018). For instance, by combining live-cell imaging and stochastic modeling, Molina et al. investigated the impact of various stimuli including TGFβ on gene expression kinetics at the CTGF locus (Molina *et al*., 2013). Similarly, Frick et al., and Tidin et al. studied the importance of relative changes (fold-changes) in the SMAD signal for the gene expression of SNAI1 and CTGF target genes at the single cell level (Frick *et al*., 2017), (Tidin *et al*., 2019). Extending on a larger set of ∼12 prototypical target genes, Lucarelli et al. demonstrated that differential SMAD complex formation is predictive for the regulation of downstream target gene expression in hepatocytes (Lucarelli *et al*., 2018). Despite these insights into SMAD signaling and gene expression, we are currently lacking quantitative, systems-level insights into the relationship between SMAD signaling dynamics, global gene expression and cell fate decisions.

In this work, we obtained a comprehensive view of how the TGFβ-induced gene-regulatory network governs cell fate decisions in mammary epithelial MCF10A cells, a well-established cellular model for EMT regulation by TGFβ. We experimentally characterized the response at multiple levels, including live-cell imaging of SMAD signaling, time-resolved genome-wide RNA sequencing and EMT assays. Using mathematical modelling, we classified target genes into potential direct targets of SMAD TFs, for which SMAD may serve as a sole input, and into indirect targets, for which regulation comprised the activity of other TFs in complex feedback and feedforward interaction. Interestingly, target genes belonging to the latter group frequently show a strong delay in gene expression, discriminate between TGFβ doses and are highly relevant for cell fate decisions such as EMT. Using TF knockdowns (KDs), we identify regulation mechanisms controlling delayed target genes: SNAI1, SNAI2, SKIL, SKI and RUNX1 function as early repressors of SMAD-dependent gene regulation, preventing premature activation, whereas JUNB serves as a delayed activator, boosting expression at late time points. We confirm that JUNB is an important enhancer of delayed gene expression upon strong stimulation, thereby specifically inducing EMT at high but not low TGFβ doses. Taken together, we employ a genome-wide systems biology approach to determine key regulatory candidates involved in SMAD signal decoding, and cell fate decisions.

## Results

### Signaling and gene expression is dose-dependent in response to TGFβ

To characterize TGFβ signaling and downstream gene expression, we used a previously established MCF10A reporter cell line with histone H2B cyan fluorescent protein tag as nuclear marker (H2B-CFP), and SMAD2 yellow fluorescent protein tag (SMAD2-YFP). As observed in earlier live-cell time-lapse imaging experiments, SMAD2 resides in the cytosol in the absence of stimulation and shows nuclear translocation upon stimulation with low (2.5 pM) and high (100 pM) doses of TGFβ (Strasen *et al*., 2018). Treatment with 2.5 pM results in a lower early signal amplitude 60 min after stimulation when compared to 100 pM. Moreover, the signal is sustained for the high dose, whereas it is transient for the low dose, reaching base line approximately 720 min post-stimulation **(Figure 1, A**).

**Figure 1:**
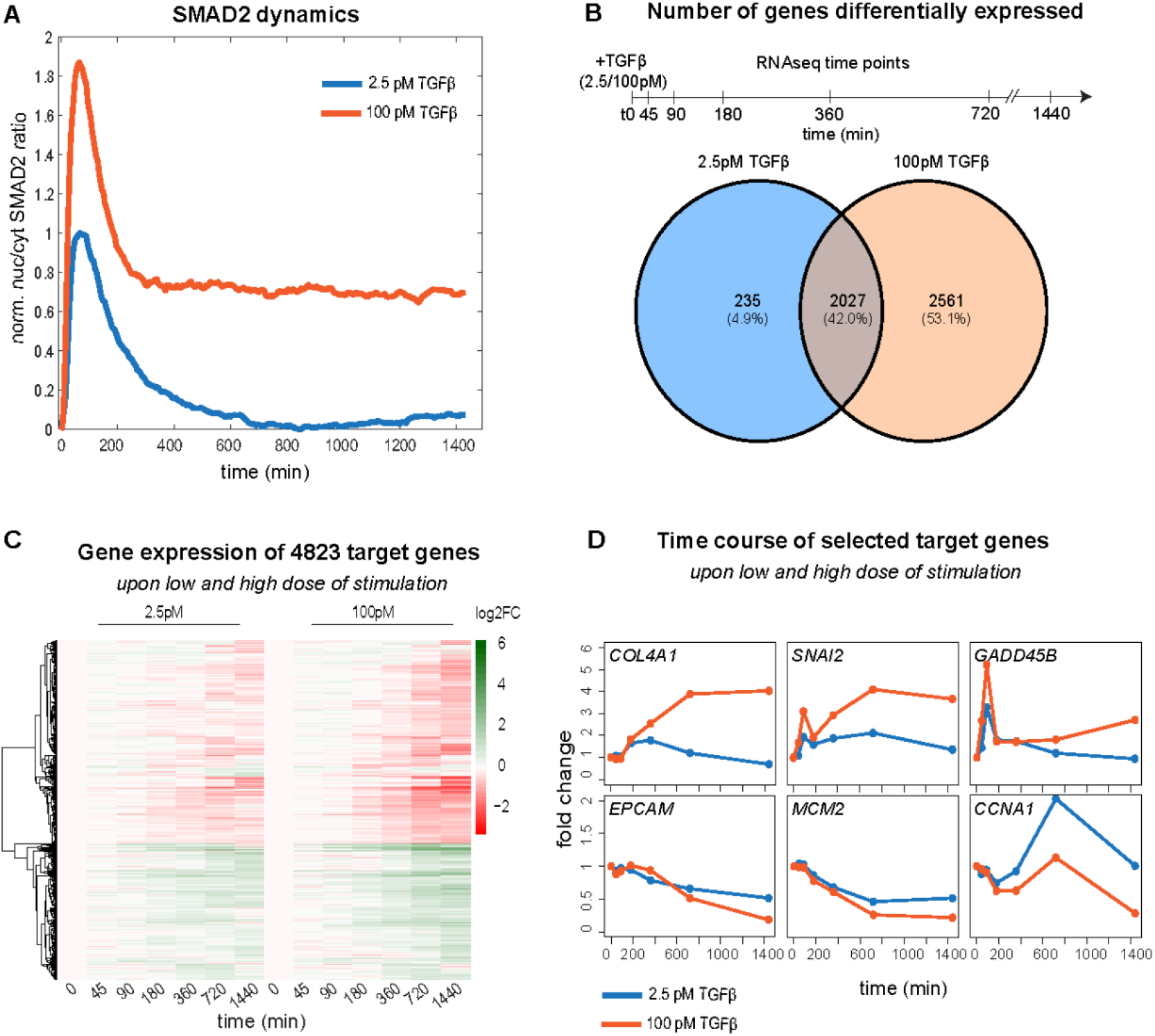
Dose-dependent signaling and gene expression in response to TGFβ. **A)** Time course of normalized nuc/cyt SMAD2 ratio upon stimulation with a low (2.5 pM, blue) and high (100 pM, orange) TGFβ dose. Shown is a background-subtracted mean of the cell population derived from the imaging data in Strasen et al. 2018 (Strasen *et al*., 2018) of 378 cells (100 pM), and 351 cells (2.5 pM). **B)** Venn diagram shows number of genes significantly (abs. FC > 1.5, adjusted p-value < 0.01) differentially expressed upon 2.5 pM and 100 pM TGFβ stimulation in MCF10A cells, in any of the time points of the sketched RNA sequencing experiment. In total, 4823 genes show altered expression compared to unstimulated control, out of which 235 (∼5 %) and 2561 (∼53 %) specifically respond at 2.5 pM and 100 pM, respectively, whereas 2027 (∼42 %) change expression in both doses. **C)** Heatmap shows trajectories of all 4823 differentially expressed genes upon low and high dose, genes (y-axis) being sorted by hierarchical clustering. **D)** Time course of known TGFβ target genes with biological functions related to EMT *(COL4A1, SNAI2, EPCAM)* and cell cycle *(GADD45B, MCM2, CCNA1)*.

Under the same stimulation conditions, we performed bulk RNA sequencing at various time points (45/ 90/ 180/ 360/ 720/ 1440 min) post-stimulation to assess global changes in gene expression. In total, 4823 genes were differentially regulated in at least one TGFβ dose and time point (p-value < 0.01, multiple testing corrected) **(Figure 1, B**). Approximately one third of the genes were upregulated by TGFβ, whereas the remainder was downregulated **(Figure 1, C**). As expected, the vast majority (2027 genes, 90 %) of the 2262 genes responding at the low dose were also differentially expressed upon strong stimulation. On top, the 100 pM stimulus specifically regulated a large set of 2561 additional genes that did not show significant changes upon treatment with 2.5 pM **(Figure 1, B)**. Hence, the higher the TGFβ stimulus, the more genes are significantly regulated. In addition, the magnitude of mRNA up- or downregulation increases in a dose-dependent gradual manner, as indicated for known TGFβ target genes **(Figure 1, D)** and in a heatmap showing the trajectories of all 4823 differentially expressed genes **(Figure 1, C)**.

### Late target genes specifically respond at high TGFβ doses

Given this dose-dependence, the question arises whether the same set of genes is regulated by both TGFβ doses or whether a subset of genes is specifically controlled upon sufficiently strong stimulation. To answer this question, we related log2-fold changes of the low and high doses at each time point, considering all genes that are differentially regulated in at least one of the two conditions **(Figure 2, A, Supplementary Figure 1, A)**. As reported in our earlier work (Bohn *et al*., 2023), we find little specificity of TGFβ-induced gene expression at early time points (90/ 180/ 360 min), as essentially the same genes are induced by the low and the high dose, with a global trend showing larger up-regulation at 100 pM TGFβ **(Figure 2, A, solid line)**.

**Figure 2:**
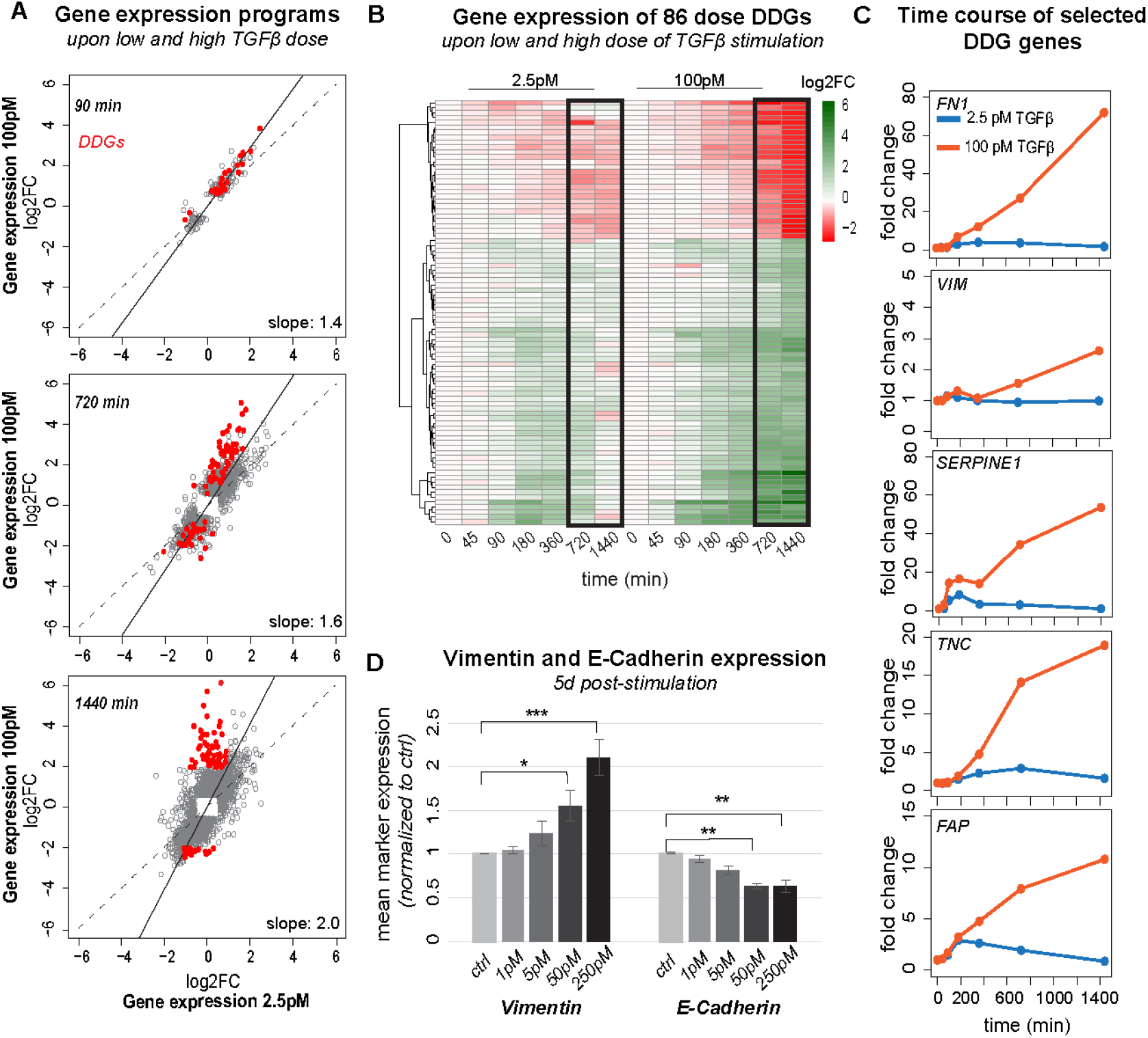
Dose-discriminating genes (DDGs) specifically respond at high TGFβ doses at late time points and are related to EMT. **A)** Scatter plot relating global gene expression changes upon 2.5 and 100 pM TGFβ stimulation at different time points (90/ 720/ 1440 min) in MCF10A cells. Each grey dot represents a significant differentially expressed gene in at least one of the two stimulation conditions, and the black solid line is a linear fit to the data. Red dots indicate DDGs differentially expressed upon high dose stimulation (absolute (abs.) fold-change (FC) > 4) but not low dose stimulation (abs. FC < 2) at late time points (720 or 1440 min post-stimulation). Number of genes: 90 min: DDGs: 24, all: 187, 720 min: DDGs: 84, all: 3009, 1440 min: DDGs: 86, all: 3920. **B)** Heatmap shows trajectories of the set of 86 DDGs defined in A. **C)** Time course of selected DDGs (*FN1, VIM, SERPINE1, TNC, FAP*) upon low and high dose TGFβ stimulation. **D)** EMT is specifically induced at high TGFβ doses. Protein levels of Vimentin and E-Cadherin measured by flow cytometry upon 5 d stimulation with different TGFβ doses (1/ 5/ 50/ 250 pM) normalized to unstimulated control. Bars comprises different number of biological replicates (n=3 for 5 pM, n=6 for ctrl, 1/ 50/ 250 pM), errors indicate SEM and significance level relative to unstimulated control (*p<=0.05, **p<=0.01, ***p<=0.001).

Similarly, at the late 720 and 1440 min time points, the vast majority of genes still shows coherent regulation by the two doses, closely following the global trend line. However, a subset of TGFβ-induced mRNAs shows atypical behavior, being strongly (up to 64-fold) upregulated upon 100 pM TGFβ stimulation, while showing essentially no response to the 2.5 pM stimulus. To comprehensively characterize these late dose-discriminating genes (DDGs), we applied filtering to the dataset, requiring that a gene shows a small fold-change (< 2-fold) at the low dose, while showing strong regulation (> 4-fold) at the high dose 1440 min after stimulation. This yielded a set of 86 DDGs, which we subjected to further analysis (**Figure 2, B, C)**.

Interestingly, the DDGs were associated with the regulation of EMT, and therefore significantly overlapped with a published EMT gene set (Fishers exact test, p-value<2·10^-16^) (**Supplementary Table 1**). Specific examples of DDGs are well-known TGFβ-induced EMT genes such as FN1, SERPINE1 and VIM, but also the cell cycle regulator TNC **(Figure 2, C)**. Gene-set-enrichment-analysis (GSEA) of DDGs indicated an overrepresentation of genes involved in extracellular-matrix remodeling, focal adhesion formation, and regulation of proteoglycans, processes associated with increased cell movement during EMT **(Supplementary Table 2)**. To validate that EMT is specifically or at least more strongly induced by the high TGFβ dose, we conducted a well-established flow cytometry-based EMT assay, measuring the up- and downregulation of Vimentin and E-Cadherin protein levels, respectively. In line with our hypothesis, the flow-cytometry data shows that only higher doses of TGFβ (50-250 pM) significantly induce EMT of MCF10A cells, whereas lower doses of 1-5 pM TGFβ cannot initiate the mesenchymal phenotype (Vimentin ^high^, E-Cadherin ^low^) **(Figure 2, D)**.

Inspection of the time courses of dose-discriminating EMT and cell cycle genes (FN1, SERPINE1, VIM, TNC) revealed exceptionally high fold-changes 720 and 1440 min after 100 pM TGFβ stimulation, with much lower or no mRNA induction in the first few hours. Such a pronounced induction delay is particularly observed for FN1, VIM, TNC, FAP, and to some extent also for SERPINE1, which shows a transient upregulation at early time points, and a more pronounced second rise 720 and 1440 min after stimulation **(Figure 2, C)**. Thus, DDGs show a late expression boost with high fold-changes, while they return to basal within 1440 min upon stimulation with the low dose. Taken together, a subset of EMT and cell cycle relevant targets shows a highly distinct expression profile for varying TGFβ doses late after stimulation, which may boost these cell fates, especially upon strong stimulation.

### Dynamics of most genes are described by a simple gene expression model

Since the temporal dynamics of gene expression are shaped by many aspects including mRNA half-life, promoter saturation and the interplay of SMADs with other TFs, we turned to mathematical modeling for a quantitative description of mRNA time courses on a genome-wide scale. In our basic gene expression model, we describe the mRNA time course using a single ordinary differential equation (ODE) which takes into account rates of mRNA synthesis and degradation **(Figure 3, A)**. The mRNA synthesis rate is assumed to depend on the nuclear SMAD2 activity (SMAD) in a sigmoidal manner (Hill equation), and the model additionally considers basal, SMAD-independent transcription (*β*_0_). As SMAD TFs may induce or repress target gene expression, we considered two variants of this model, an activator (1) and an inhibitor (2) model

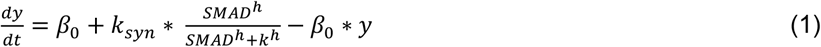

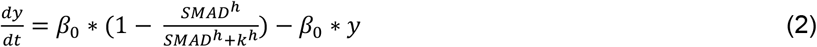

where *y* is the fold-change in target gene expression, *β*_0_ the basal transcription and degradation rate, *k*_*syn*_ the maximal transcription rate, SMAD the SMAD2 input signal, *h* the Hill coefficient, and *k* the concentration of promoter half-saturation.

**Figure 3:**
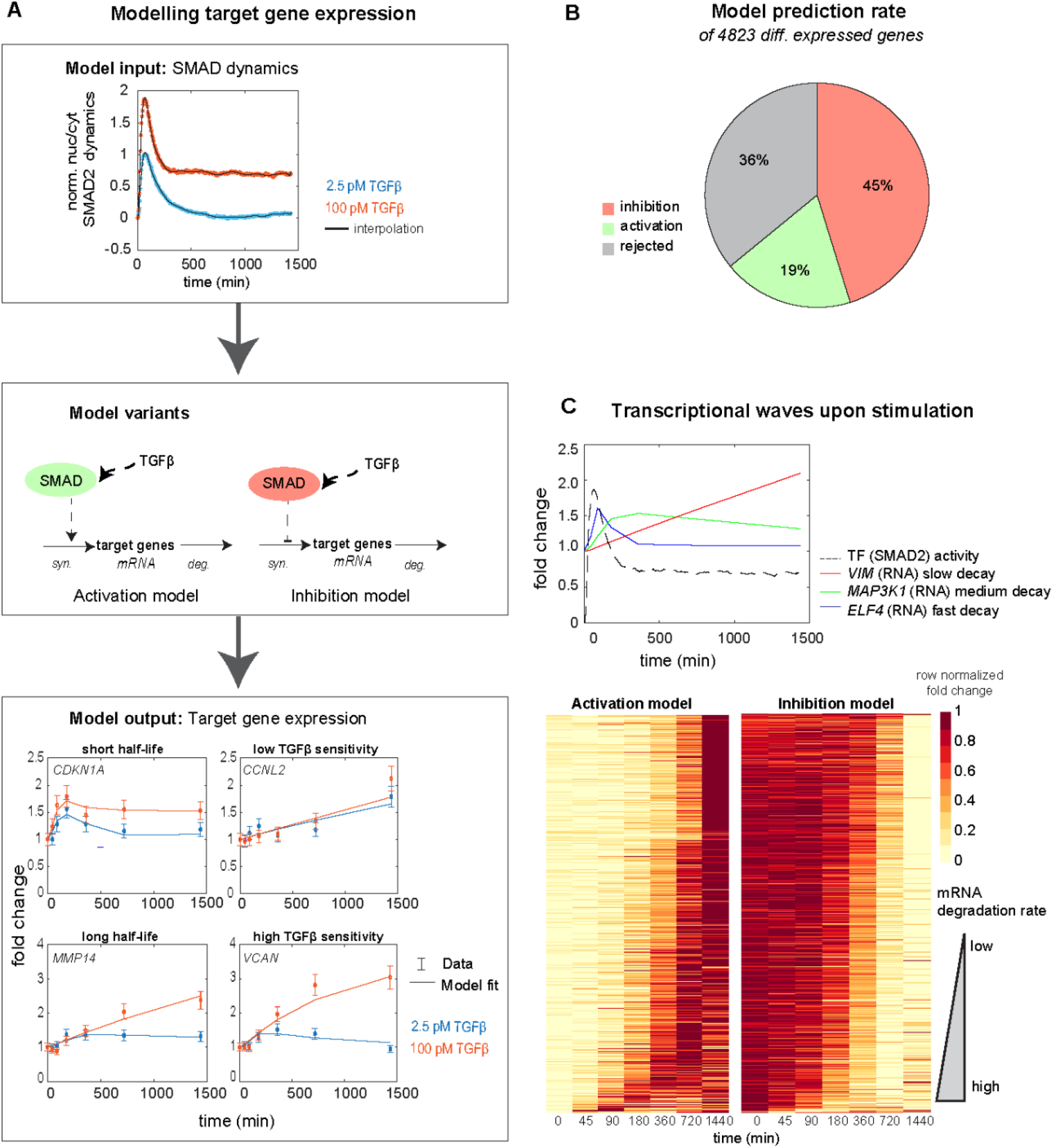
Modeling SMAD-dependent activation and repression of gene expression. **A)** Modeling framework: experimentally measured SMAD2 dynamics (top) serve as model input, controlling target gene transcription in an ODE describing mRNA synthesis and degradation (middle). In two model variants, target gene activation and repression by SMAD are considered, and the simulated RNA time course is fitted to the RNA sequencing data (bottom). The ODE-model describes target genes with short (*CDKN1A*) and long half-lives (*MMP14*), and by adjusting the Hill equation parameters genes with low (*CCNL2*) and high (*VCAN*) TGFβ sensitivity. **B)** Proportions of explained and rejected target genes. 64 % (3091 genes) of 4823 target genes can be explained by either activation (19 %, 919 genes) or inhibition (45 %, 2160 genes) model, whereas 36 % (1744 genes) are rejected as judged by χ^2^-testing of model fitting results (see Methods). **C)** Model predicts the order of target gene expression based on mRNA degradation rates. Top: Simulated time courses using the best-fit parameters of three exemplary genes, one with fast degradation (*ELF4*, blue), one with medium degradation (*MAP3K1*, green), and one with slow degradation (*VIM*, red). Bottom: Heatmap showing measured time courses of target genes explained by the activation (919 genes) and inhibition (2160 genes) models after sorting according to the best-fit mRNA degradation rate. Each row is min-max normalized by its minimal and maximal expression values.

To assess whether these simple models agree with our data, we used the measured SMAD2 nuclear translocation as a model input, and fitted the simulated mRNA fold-change to the corresponding RNA sequencing measurements of each gene, adjusting Hill equation parameters, and the mRNA degradation rate (β0). For each of the 4823 differentially expressed genes, the activator and inhibitor models were tested separately, simultaneously considering the nuclear-to-cytoplasmic (nuc/cyt) SMAD2 ratio, and RNA sequencing data at low and high TGFβ doses **(Figure 3, A)**. Based on the fitting results, we concluded that 64 % of all differentially expressed target genes could be quantitatively explained by one of the two models, 19 % and 45 % being described by the activator and inhibitor version, respectively (Chi2-test, see Methods) **(Figure 3, B**). Taken together, minimal SMAD2-dependent mRNA expression models account for a large part of the global gene expression program, suggesting that much of dose-dependence simply follows gradual changes in SMAD2 activity and does not require additional input by other signaling pathways or TFs.

The dynamics of the explained target genes show a large variation, some mRNAs reacting fast whereas others do so only with a delay **(Figure 3, A, C)**. Our model accommodated this different kinetics by adjusting mRNA half-life: genes with a short, fitted mRNA half-life rapidly follow the transient SMAD2 increase and decline (e.g., CDKN1A), whereas long-lived mRNAs show a late response and remain differentially expressed long after the SMAD2 input has vanished (e.g., MMP14) **(Figure 3, A)**. Hence, in line with previous reports (Hao and Baltimore, 2009), the mRNA degradation rate is the major controller of the temporal order of gene expression **(Figure 3, C)**. In contrast, the Hill equation parameters primarily determine TGFβ dose sensitivity, as some mRNA show a strong difference between the low and the high TGFβ doses (e.g., CDKN1A, MMP14, VCAN), while others show saturation already at the low TGFβ dose (e.g., CCNL2) **(Figure 3, A)**. Taken together, our modeling framework classifies genes according to mechanisms of regulation and provides mechanistic explanations for the dynamics of the explained genes.

### Genes with complex regulation show signs of feedforward regulation

Out of 4823 differentially expressed genes in response to TGFβ stimulation, 1744 (36 %) could not be explained by the simple SMAD-dependent ODE-models and therefore show signs of complex regulation. Many of these 1744 rejected genes seem to be important for cellular decision making, as they contain 156 of 391 differentially regulated cell cycle genes and 93 out of the 172 differentially regulated EMT genes **(Supplementary Table 3)**. Manual inspection of time courses identified five recurrent time course patterns **(Figure 4, A, B):**

**Figure 4:**
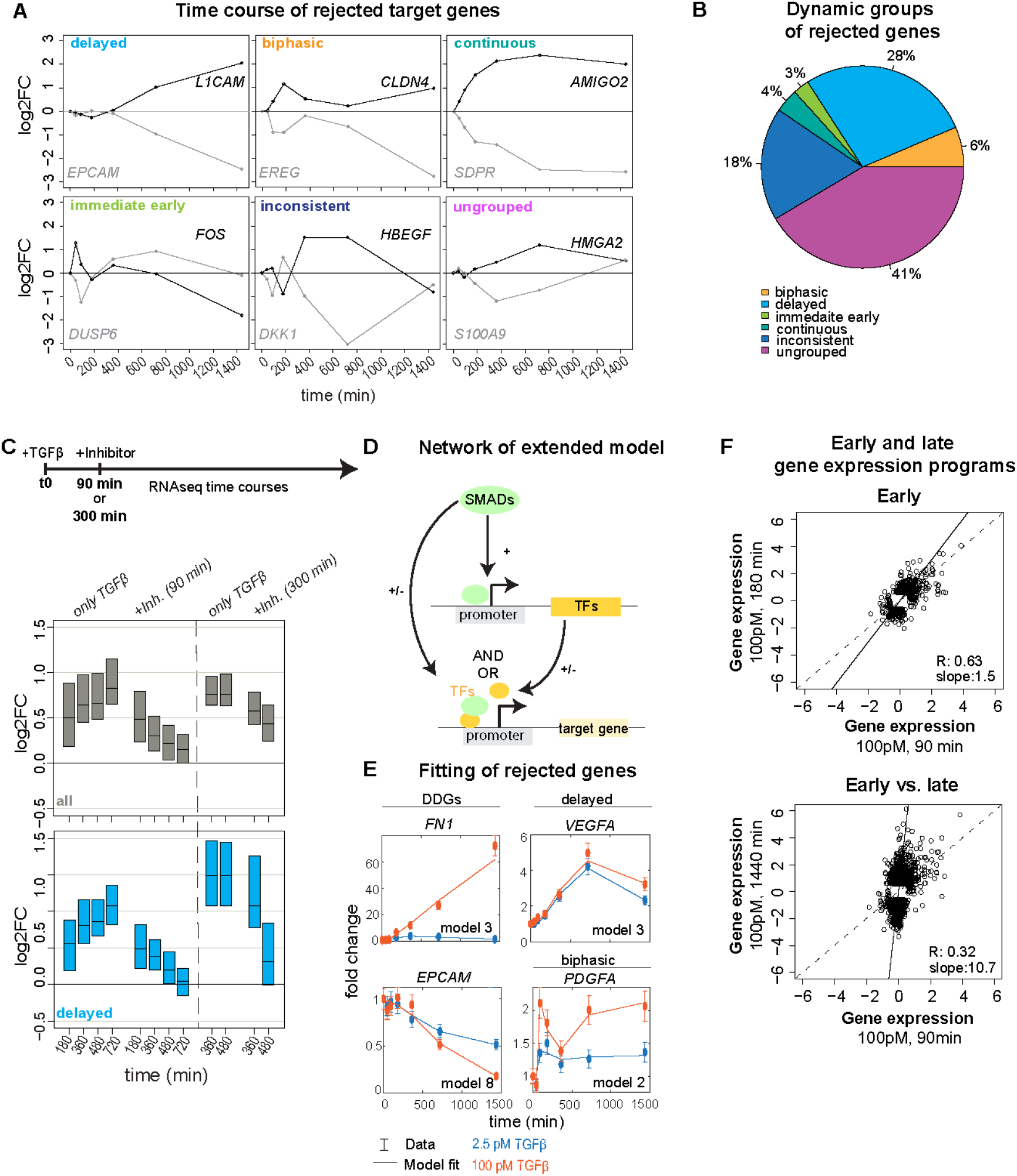
Evidence for feedforward regulation in SMAD-dependent gene expression. **A)** RNA sequencing time courses (100 pM TGFβ) of selected genes rejected by the kinetic models in Figure 3, gene names being indicated next to each trajectory. The following temporal dynamics can be observed: delayed (late onset of differential expression), biphasic (fast and sustained differential expression with intermediate decline), continuous (fast and sustained differential expression), immediate early (early differential expression followed by rapid decline), and inconsistent kinetics (early up- and late downregulation, or vice versa). Ungrouped genes like *S100A9* and *HMGA2* show distinct dynamics and/or low expression changes, and can therefore not be assigned to any of the dynamic groups. **B)** Automated classification of all 1744 rejected genes into the five time course patterns (immediate early, biphasic, continuous, delayed and inconsistent kinetics) using expression cutoffs at multiple time points (see Methods and Supplementary Table S3). **C)** RNA sequencing after TGFβ-receptor inhibitor treatment confirms ongoing SMAD-dependency of late target genes. Cells were stimulated with 100 pM TGFβ at t=0 and the TGFβ receptor inhibitor SB431542 was applied at 90 or 300 min. Genome-wide RNA sequencing was performed at the indicated time points, either in cells treated with TGFβ only (left) or after inhibitor addition (right). The expression distributions (boxes: median with lower and upper quartile) of all upregulated (log2FC > 0.58, adj. p-value < 0.01) TGFβ target genes (top, grey, + 90 min: 789 genes, + 300 min: 993 genes) or those belonging to the delayed gene expression group (bottom, blue, + 90 min: 112 genes, + 300 min: 88 genes) declines over time, upon inhibitor treatment. See SF3 for analysis of downregulated genes and other gene groups. **D)** Schematic representation of extended feedforward loop (FFL) model. In the FFL model, SMAD upregulates (+) the expression of a hypothetical TF (TF), which in turn regulates target genes jointly with SMAD. Both SMAD and the TF function either as a repressor (−) or activator (+), and control on the target gene promoter with AND- or OR-gate logic, giving rise to in total 8 model variants (see methods for details and supplementary Figure 3, G for model variants, and Supplementary Table 5 for FFL fitting results). **E)** Time courses fits of selected DDGs (*FN1, EPCAM*), delayed (*VEGFA*), and biphasic (*PDGFA*) target genes with optimal extended model version indicated on the bottom right. **F)** Distinct early and late gene expression programs induced by TGFβ. Scatter plot relating log2 fold-changes relative to unstimulated control of two early time points (top, 90 min vs. 180 min) or early vs. late time points (bottom, 90 min vs. 1440 min). Each dot represents a significant differentially expressed gene in at least one stimulation condition (top: 516 genes, bottom: 3722 genes). Lines: bisecting (dashed) and linear fit to the data (solid), slope and correlation coefficient indicated on the bottom right.

I. delayed kinetics: strong rise, but with a pronounced delay
II. inconsistent kinetics: early up-followed by late downregulation (or vice versa)
III. biphasic kinetics: transient expression followed by a second, late rise (or drop)
IV. continuous kinetics: continuous up-or downregulation despite a transient SMAD input
V. immediate early kinetics: early upregulation followed by rapid decline (‘spike like’)

By time course filtering, **(see Methods and Supplementary Table 4)** we could assign a large number of 1021 out of 1744 rejected genes to one of these five dynamic gene groups, the majority belonging to the delayed (483), inconsistent (315) and biphasic (112) classes **(Figure 4, A, B, Supplementary Figure 2 A-E, Supplementary Table 4)**. The remaining 723 out of 1744 rejected genes, not assigned to any of the dynamic gene groups (i-v), tend to show minor induction or repression by TGFβ (< 2.5-fold), especially at late time points **(Supplementary Figure 2, F**). Taken together, many genes rejected by our simple kinetic models show signs of late expression regulation after the SMAD signal has reached plateau (delayed, biphasic, inconsistent, and continuous groups). In line with late gene expression regulation being important for TGFβ dose-discrimination **(Figure 2)** the 55 DDGs that could not be described by the SMAD-dependent gene expression models were classified into delayed (33 out of 55), continuous (21 out of 55) and biphasic (1 out of 55) classes **(Supplementary Figure 1 B, C)**.

Previous studies suggested that late target gene regulation by TGFβ involves feedforward loops (FFLs), in which the input (SMAD2) regulates late target genes by a combination of direct and indirect mechanisms, i.e., direct SMAD binding to the gene promoter and by controlling the expression of transcriptional co-regulators (Sundqvist *et al*., 2018). In line with such feedforward regulation, many early target genes expressed within the first 90 min upon 100 pM TGFβ stimulation encode for TFs and epigenetic regulators (39 out of 109 differentially expressed genes, 39 %) **(Supplementary Figure 3, A, B)**. To test whether late target genes still directly depend on the initial SMAD signal, we treated cells 90 or 300 min after 100 pM TGFβ stimulation with SB431542, a small molecule inhibitor of TGFβ receptors, which quickly downregulates SMAD signaling with a half-life of ∼60 min (Inman *et al*., 2002), (Strasen *et al*., 2018). Following gene expression over time using RNA sequencing, we observed a quick downregulation of almost all TGFβ target genes within 60-120 min of SB431542 application when compared to cells that had not been treated with the inhibitor. This suggests a continuing dependence on the SMAD signal, which importantly was also observed for late target gene sets belonging to the classes with delayed, continuous or biphasic kinetics, and for those showing dose-discrimination **(Figure 4, C, Supplementary Figure 3, D, E, F)**.

To further validate large-scale feedforward regulation of the 1744 rejected target genes, we extended our kinetic models of gene expression by a hypothetical SMAD-regulated TF **(Figure 4, D)**. This TF is described by the synthesis-degradation model (see methods), and in turn regulates the synthesis of the target gene of interest. For model fitting, we again employed the SMAD input and target gene measurements at low and high TGFβ doses, while leaving the dynamics of the unknown TF unconstrained. Since both SMAD2 and TF may function as activators or repressors of gene expression, and they may interact on target gene promoters with AND- or OR-logic, we considered in total 8 model variants **(Supplementary Figure 3, G)**. After fitting and statistical testing, we found that 1577 out of 1744 initially rejected target genes could be quantitatively explained by the extended FFL models **(Figure 4, E, Supplementary Table 5)**. Of these, 1074 out of 1577 (68 %) were explained with the co-TF acting as an activator, while for 503 out of 1577 (32 %) target genes the co-TF functioned as a repressor. When combined, the minimal model and its FFL derivatives explained the expression of 4656 out of 4823 (96.5 %) SMAD target genes, suggesting that the late TGFβ-induced gene expression program is shaped by both repressors and activators of target genes.

As feedforward regulation may lead to the upregulation of distinct genes late after stimulation, we tested whether gene expression is reprogrammed over time. To this end, we globally related fold-changes in target gene expression across time points for the 100 pM TGFβ stimulus, and indeed found highly distinct gene expression programs between early (90 min) and a late (720/ 1440 min) time points, many genes showing specific expression late after stimulation. In contrast, the response at two early time points (90 and 180 min) were highly similar **(Figure 4, F, Supplementary Figure 3, C)**. Moreover, even though TFs and epigenetic regulators were also induced early upon 2.5 pM TGFβ stimulation (22 out of 53 diff. reg. genes, 41 %) **(Supplementary Figure 3, A, B)** their average fold-change was significantly lower compared to 100 pM TGFβ. Consistently, the early and late gene expression programs at 90 and 720 min stimulation were highly similar at the low dose **(Supplementary Figure 3, C)**, confirming specific late transcriptional reprogramming at high TGFβ doses.

### Feedforward regulation by early repressors and JUNB as a late activator determines DDG expression

To identify potential regulators of late gene expression, we turned to TF KDs followed by global RNA sequencing. For the identification of candidates, we used a curated list of TFs and epigenetic regulators (Lambert *et al*., 2018), and focused on those induced within the first 90 min after TGFβ stimulation, thereby obtaining a shortlist of 7 co-factors (SNAI2, JUN, JUNB, KLF10, SKIL, RUNX1, SKI) **(Figure 5, A)**. Additionally, we included SMAD2 and SMAD3 KDs as positive controls, and furthermore considered SNAI1 and ATF3, which are known induced TFs upon TGFβ stimulation but did not reach significance in our data set (Kang, Chen and Massagué, 2003; Yin *et al*., 2010; Zhang *et al*., 2014).

**Figure 5:**
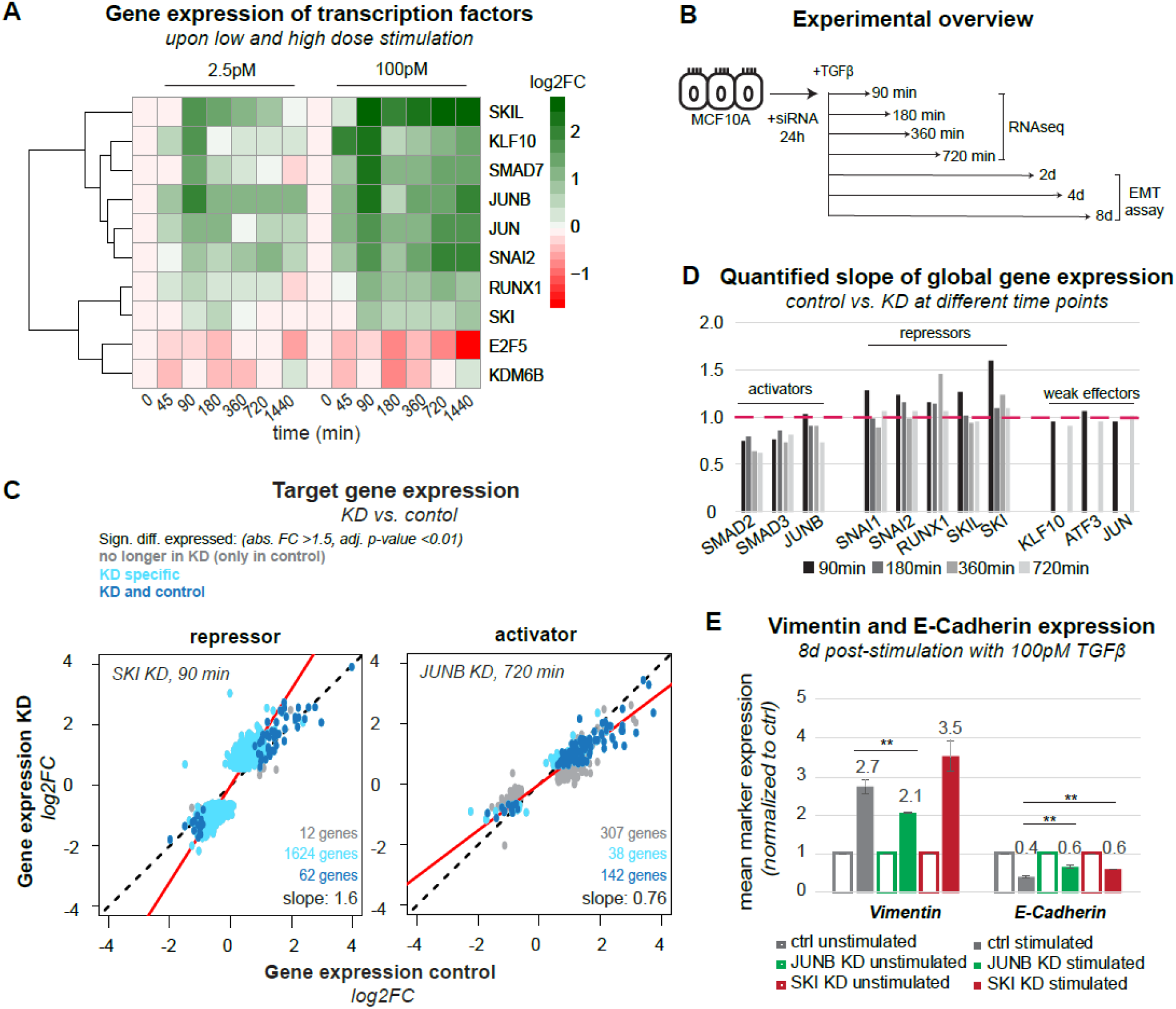
TFs shaping TGFβ target gene expression and cell fates. **A)** Gene expression time course of TFs (*SKIL, KLF10, SMAD7, JUNB, JUN, SNAI2, RUNX1, SKI, E2F5*), and epigenetic regulators (*KDM5B*) with differential expression (adj. p-value < 0.01, abs. F C> 1.5) in the first 90 min after TGFβ stimulation. **B)** Experimental set up showing time points of RNA sequencing (90, 180, 360, 720 min) and EMT assays (2, 4, 8 d) post-stimulation with 100 pM TGFβ of MCF10A cells pre-incubated with siRNA targeting TFs for 24 h. **C+D)** KD reveals activators and repressors of SMAD-mediated gene regulation. **C)** Scatter plot showing TGFβ-induced global gene expression changes in control vs. SKI (90 min, left) and control vs. JUNB (720 min, right) KD. The slope of a linear trend line (red line) across all genes determines the strength of the repressor (left) or activator (right) effect. Genes significantly differentially expressed (adj. p-value < 0.01, abs. FC > 1.5) in control only, KD only or control and KD are colored in grey, blue, and turquoise, respectively. **D)** Quantified slope of KD effect on global gene expression, as described in C, for the complete set of tested TFs and all time points (90, 180, 360, 720 min). SMAD2 and SMAD3 are activators at all time points, whereas JUNB is a late activator; repressors SNAI1, SNAI2, RUNX1, SKIL, SKI mainly act at early time points. KLF10, ATF3, and JUN show weak effects. **E)** Flow cytometry analysis of epithelial (E-Cadherin) and mesenchymal (Vimentin) marker expression upon long-term (8 d) TGFβ stimulation of control and KD (SKI and JUNB KD) MCF10A cells, n=3, SEM is shown (**p<=0.01). JUNB KD represses and SKI KD enhances the mesenchymal phenotype.

For KD experiments, cells were pre-incubated with siRNAs for 24 h and then subjected to stimulation with a saturating concentration of TGFβ (100 pM) **(Figure 5, B)**. Samples were taken at the time point of stimulation (t0) or 90, 180, 360, and 720 min after stimulation. For each of the KDs, we validated highly efficient depletion of the targeted TF **(Supplementary Figure 5, A)** and performed bulk RNA sequencing to determine global changes in gene expression. The KD of SMAD2, SNAI1, SNAI2, and SKI had pronounced effects on basal gene expression before stimulation, with 485, 846, 1254, and 268 genes showing an absolute fold-change greater than 1.5, respectively, whereas basal effects were negligible for the remaining factors (JUNB, SKIL, RUNX1, SMAD3, JUN, KLF10, and ATF3) **(Supplementary Figure 5, B)**.

To quantify KD effects on TGFβ-induced gene expression, we related the log2-fold changes induced by TGFβ stimulation in control vs. KD cells across all genes **(Figure 5, C, D)**. In general, we found a strong correlation of TGFβ-induced log2-fold changes (R = 0.87 – 0.99), suggesting that, in most cases, a similar overall gene expression program was activated by TGFβ in control and KD cells **(Supplementary Table 6)**. By fitting a linear trend line to the data, we determined the direction of change and the magnitude of the KD effect on global target gene expression **(Figure 5, C, solid red line)**. TFs with slopes > 1 upon KD were considered as repressors, as their depletion leads to target gene upregulation, whereas the opposite (slope < 1) is true for activators. Based on this criterion and statistical analysis, KDs of SNAI1, SNAI2, JUNB, SKI, SKIL, and RUNX1 as well as SMAD2, and SMAD3 had significant effects on global TGFβ-induced gene expression (t-test, α=0.05). We found that RUNX1, SKIL, SKI, SNAI1, and SNAI2 function as repressors (slope >1), whereas JUNB, SMAD2 and SMAD3 function as activators (slope <1) **(Figure 5, C, D Supplementary Table 6)**.

Given the KD analysis at different time points, we could determine the timing of TF action on target gene expression: As a general trend, the repressors primarily acted shortly after TGFβ stimulation, with SKI showing the strongest effect on target gene expression at 90 min among all repressors. Within the activators of TGFβ target gene expression, SMAD2 and SMAD3 had pronounced effects at all time points (90/ 180/ 360/ 720 min) as expected, whereas JUNB specifically acted at the late time point, consistent with the reported role of JUNB being a feedforward regulator of SMAD-induced gene expression that redirects SMADs to novel target genes (Sundqvist *et al*., 2018). To exclude that changes in gene expression upon TF KD arise from feedback to the SMAD signaling pathway, we examined the nuc/cyt translocation of SMAD2 upon KD of JUNB, SKIL, and SKI **(Supplementary Figure 5, C, D, Supplementary Table 7)**. We observed only slight alterations in SMAD2 translocation, excluding that feedback regulation is the major cause of changes in gene expression.

Some TFs tended to globally affect all target genes, whereas others have the most pronounced effects on a subset of target genes, therefore controlling a specific gene expression program **(Figure 5, C, Supplementary Figure 6)**. To quantify such TF specificity, we determined the number of genes that show a significantly different TGFβ response in the KD when compared to non-targeting control. For some KDs the TGFβ response of almost all target genes is homogeneously affected (SMAD2, SMAD3) **(Supplementary Figure 6)**, whereas for other KDs only a subset of genes shows a differential TGFβ response, and is observed for early repressors (SKIL, SKI, RUNX1, SNAI1 and SNAI2, 90-360 min) and JUNB as a late activator (720 min). **(Figure 5, C, Supplementary Figure 6**, compare dark blue, light blue and grey dots).

Taken together, the TFs decompose into repressors of SMAD-induced gene expression primarily acting at early time points (SKIL, SKI, RUNX1, SNAI1, SNAI2), and into activators that are either general mediators of the signal at all time points (SMAD2, SMAD3) or specifically act late (JUNB), possibly as feedforward amplifiers. Six factors seem to control a specific subset of SMAD target genes (SKI, SKIL, SNAI1, SNAI2, RUNX1, JUNB), indicating that they may reshape the SMAD-induced gene expression program. The remaining factors JUN, KLF10 and ATF3 had a weak impact on overall gene expression patterns, both before and after TGFβ stimulation **(Supplementary Figure 5, B, 6)**.

### SMAD cofactors modulate TGFβ-induced EMT

To confirm that regulators of late gene expression affect cellular decision making, we quantified EMT by measuring protein levels of Vimentin and E-Cadherin upon KD of a co-activator (JUNB) and a co-repressor (SKI) 2-8 days after stimulation with 100 pM TGFβ. To maintain the KD and to replenish TGFβ in these long-term experiments, we re-treated the culture with fresh siRNAs and TGFβ every 2 d, and confirmed that the downregulation of the targeted TF was stable over time **(Supplementary Figure 7, A)**.

Compared to unstimulated control, TGFβ-induced EMT induction is weakened in JUNB KD cells, as evidenced by less pronounced Vimentin upregulation and E-Cadherin downregulation 2, 4 **(Supplementary Figure 7, B, C)** and especially 8 d post-stimulation **(Figure 5, E)**. This decreased sensitivity to EMT induction by TGFβ is consistent with JUNB being an activator of late TGFβ-induced EMT genes (see above). On the contrary and as expected, the transcriptional repressor SKI blocks EMT induction, as Vimentin was more upregulated by TGFβ in SKI KD compared to control, 8 d after stimulation **(Figure 5, E)**. However, at earlier time points (2 d and 4 d post-stimulation), SKI KD led to lowered Vimentin induction by TGFβ, suggesting that SKI might actually enhance the onset of EMT **(Supplementary Figure 7, B)**.

Taken together, our data places JUNB as an activator of EMT in MCF10A cells, whereas SKI may play a dual role in this process, exerting its repressive role only at late time points.

### JUNB reinforces the dose discrimination at the level of target gene expression

Given its role as an amplifier of late gene expression and EMT induction, JUNB may contribute to TGFβ dose-discrimination, specifically boosting the late DDG expression upon high dose stimulation. To confirm such late amplification, we compared time courses of selected DDGs (e.g., FN1, SERPINE1, SLC46A3, TNC) in JUNB KD and control conditions using the previously described RNA sequencing data upon 100 pM TGFβ stimulation (**Figure 6, A**). As expected, JUNB KD specifically eliminated the late DDG amplification 720 min post-stimulation (FN1, SLC46A3, TNC) or had the more pronounced effect at this late time point (SERPINE1). In contrast, KD of the constitutive regulator SMAD2 diminished expression values of most DDGs already at earlier time points (SERPINE1, SLC46A3, TNC) and depletion of the early repressor SKI either had no effect on DDG expression (FN1, SLC46A3, TNC) or led to early upregulation (SERPINE1) (**Figure 6, A**). Having shown that JUNB promotes the late amplification of specific DDGs, we asked whether the same holds true on the complete set of 55 DDGs, which are not explained by the simple gene expression model. Indeed, our analysis confirms that JUNB KD had little effects on TGFβ-induced fold changes at early time points, while leading to a homogenous and significant reduction for the 55 DDGs at 720 min **(Figure 6, B)**. In contrast, depletion of SMAD2 reduced global TGFβ-induced DDG expression already at the earliest time points, reaching significance 360 and 720 min post-stimulation **(Figure 6, B)**.

**Figure 6:**
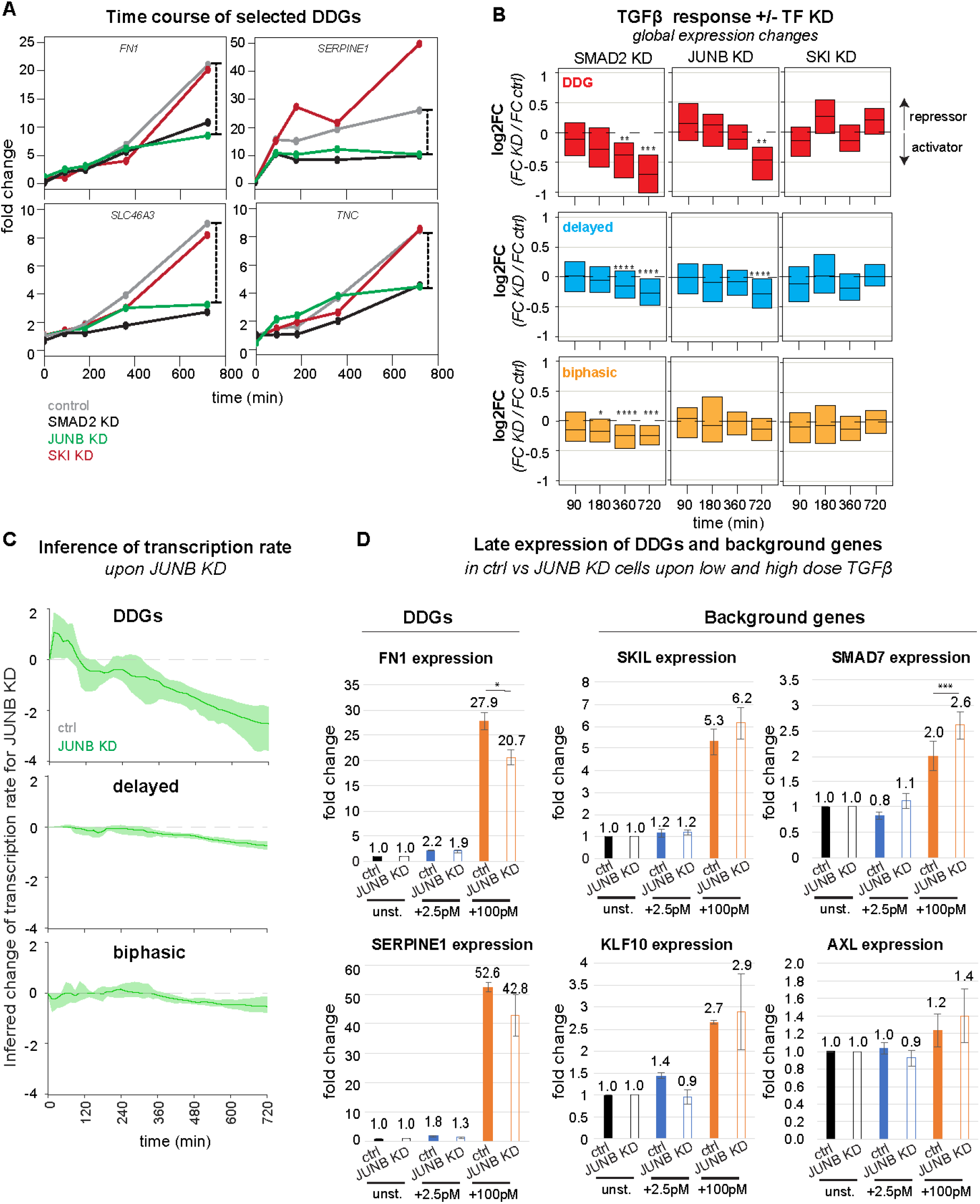
JUNB reinforces TGFβ dose-discrimination. **A)** JUNB KD boosts late expression of selected DDGs upon high dose TGFβ stimulation. Shown are time courses of DDGs (*FN1, SERPINE1, SLC46A3, TNC*), defined by late amplification for the strong TGFβ stimulus (Figure 2) in control (grey), SMAD2 KD (black), JUNB KD (green), and SKI KD (red) upon 100 pM TGFβ stimulation (same RNA sequencing data as in Figure 5, B). Scale on the right indicates expression difference between control and JUNB KD condition at 720 min. **B)** JUNB KD globally boosts late expression of DDGs (red, 43 genes), delayed target genes (blue, 290 genes), and biphasic target genes (yellow, 64 genes) rejected by the simple gene expression model (Figure 3). Boxes show changes of TGFβ-induced gene expression upon KD (log2FC = (KD at t=x/ KD at t=0) / (control at t=x/ control at t=0)) as a distribution across all genes belonging to the indicated gene set (median with lower and upper quantile). SMAD2 KD significantly reduces TGFβ-induced gene expression changes at earlier time points compared to JUNB KD (t-test). See Supplementary Figure 8, A, B for analysis of downregulated genes and other gene groups. (*p<=0.05, **p<=0.01, ***p<=0.001, ****p<=0.0001) **C)** Time-dependent target gene transcription rates (v(t)) of gene groups were inferred from RNA-sequencing data for JUNB KD and control cells. The transcription rate was separately inferred for each gene using the equation 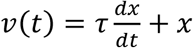, where x is the mRNA expression time course interpolated from RNA sequencing data using constrained cubic splines (details see methods). To quantify the effect of JUNB, the changes in the transcription rate upon JUNB KD (Δv = *v*_KD_ − *v*_control_) were calculated for TGFβ-upregulated DDGs (top), delayed (middle) and biphasic (bottom) genes. Lines: median of Δv in each across all genes in each group. Shades: bootstrapped confidence bands (2000 bootstrap samples). Shown are the results for TGFβ-upregulated genes; see Supplementary Figure 8, C for and the same inference of TGFβ-downregulated genes. **D)** JUNB KD reduces dose discrimination of by DDGs. MCF10A control and JUNB KD cells were stimulated with 2.5 (blue) and 100 pM (orange) TGFβ for 1440 min, and the expression of FN1 and SERPINE1 (n=3) was assessed by qPCR. Additionally, the expression of a background gene set belonging to the delayed and biphasic kinetic groups, including SKIL, SMAD7, KLF10, and AXL (n=3) was analyzed. SEM is shown.

To test whether the late amplification by JUNB is exclusive to the set of DDGs, we also analyzed the impact of SMAD2, JUNB, and SKI on other dynamic gene groups that could not be explained by the simple gene expression model **(Figure 6, B, Supplementary Figure 8 A, B)**. For the delayed gene group, which is closely related to the DDGs, we again observed a specific and significant JUNB KD effect at the late time point, which however was small compared to the DDGs. For other gene groups, JUNB KD had no significant effect on TGFβ-induced expression (biphasic and immediate early gene groups), or it barely reached significance (continuous gene group). Taken together, these data confirm the JUNB-mediated late amplification is most pronounced in DDGs when compared to other gene groups. In line with a specific late amplification by JUNB, the SMAD2 KD impacts all gene groups already at earlier time points **(Figure 6, B, Supplementary Figure 8 A, B)**.

To further confirm the late DDG amplification by JUNB, we directly inferred time-dependent transcription rates of target genes in JUNB KD vs. control conditions **(Figure 6, C)**. For inference we used a generic ODE describing mRNA expression as a function of transcription and mRNA degradation (see Equation 23), and rearranged the formula to express the transcription rate as a function of target gene mRNA slope and expression level in the RNA sequencing data (see Equation 24 and see Methods for details). As expected, the transcription rate of DDGs was strongly reduced upon JUNB KD, specifically at late time points **(Figure 6, C top)**. Delayed genes showed a similar drop in transcription, but with much milder amplitude **(Figure 6, C middle)**, and biphasic genes showed almost no (early time points) or minor (late time points) change upon JUNB KD **(Figure 6, C bottom)**. This regulatory pattern was observed in both, genes upregulated by TGFβ **(Figure 6, C)**, and genes downregulated by TGFβ **(Supplementary Figure 8, C)**. Thus, our transcription-rate inference supports that JUNB is specific regulator for late DDG amplification. To directly test whether JUNB mediates the decoding of TGFβ doses, we analyzed the expression of two EMT-relevant DDGs (SERPINE1 and FN1) in control and JUNB KD cells upon stimulation with a low (2.5 pM) and high (100 pM) TGFβ dose for 1440 min. In line with our expectation, we found that the JUNB KD specifically reduced SERPINE1 and FN1 expression in high dose conditions, with lesser effects at the low dose **(Figure 6, D)**. In contrast, a background gene set, comprising rejected target genes with delayed expression kinetics but not classified as DDGs (SMAD7, SKIL, KLF10, and AXL), did not show a specific response at the high TGFβ dose and was generally insensitive to JUNB KD **(Figure 6, D)**. Taken together, our data suggests that JUNB is involved in the late amplification of DDGs, and therefore may contribute to specific decision-making towards EMT for sufficiently strong TGFβ stimuli.

## Discussion

Understanding how ligand binding to cell surface receptors controls intracellular signaling, gene expression changes and ultimately cellular fate is critical for the development of therapeutic strategies targeting the TGFβ-pathway. In this work, we focused on specificity mechanisms in TGFβ-induced gene regulation, and characterized how TFs dynamically interact to drive changes in gene expression, and how this may be involved in shifting of SMAD signaling from a tumor suppressive to a tumor promoting function. Previous studies have demonstrated that the temporal dynamics of signaling molecules such as ERK, NFkB and p53 determine phenotypic outcomes (Purvis and Lahav, 2013). Consequently, our objective was to explore the mechanisms responsible for decoding SMAD signaling dynamics into specific gene expression programs and cell fate decisions.

In our recent work, we reported that SMAD-induced gene expression exhibits little specificity within the first few hours after stimulation, as two TGFβ family ligands, GDF11 and TGFβ, induced very similar transcriptome changes across ligand doses and cellular stages, including quiescence and proliferation (Bohn *et al*., 2023). Previous studies reported distinct early and late gene expression programs in response to TGFβ treatment (Yang *et al*., 2003; Sundqvist *et al*., 2018). Following up on this, we performed time-resolved RNA sequencing at a low and high dose TGFβ stimulation to determine whether dose-specific gene expression programs emerge at later time points. Indeed, we found a subset of targets, termed dose-discriminating genes (DDGs), that was selectively regulated upon high dose TGFβ stimulation after 12-24 h (Figure 2). Many of these DDGs show weak expression early after stimulation and are strongly amplified at late time points, suggesting a requirement for TFs that are induced by the TGFβ signal with a delay. To classify target genes into those that simply respond to the SMAD signal and those that require additional input by late TFs, we devised a genome-wide modeling approach, in which the temporal evolution of each target gene is described taking into account the balance between transcription and mRNA degradation. Considering the experimentally measured SMAD input and fitting the model parameters to describe the RNA sequencing time courses, we could explain 64 % of the 4823 differentially expressed target genes by this simple model (Figure 3). Similarly, gene expression models with an experimentally measured TF input disentangled input decoding in other cellular networks such as the p53 and MAPK signaling pathways (Purvis *et al*., 2012; Uhlitz *et al*., 2017) and were also used to describe the dynamics of a small set of 12 TGFβ target genes in liver cells (Lucarelli *et al*., 2018). Likewise, similar dynamic models were used to infer mechanisms of biological regulation from multi-OMICS time course data (Peshkin *et al*., 2015; Becker *et al*., 2018). Interestingly, even though our model considered a flexible Hill-type sigmoidal function for transcriptional regulation, it failed to describe the dynamics of ∼1700 target genes, including the majority of DDGs. This suggests that late dose-dependent amplification cannot be explained by switch-like promoter regulation but additionally requires input from dynamically changing TFs.

Following up on this, we conducted KD experiments followed by RNA sequencing to elucidate the influence of TGFβ-induced TFs on target gene expression and found pronounced time-dependent regulatory effects: Among the tested TGFβ-induced TFs, JUNB showed activator function, mainly acting at late time points, whereas the remaining co-regulators (SNAI1/2, SKI, SKIL, RUNX1) were classified as early repressors (Figure 5, D). In line with our findings, SNAI2 has been identified as a key regulator of mammary epithelial cell identity (Phillips and Kuperwasser, 2014). Even though their time-dependent effects have not been characterized in detail, SKIL and SKI are known to be repressors of the TGFβ-induced gene expression response, (Tecalco-Cruz *et al*., 2018) which control EMT in non-small cell lung cancer (Yang *et al*., 2015) and metastasis in breast cancer cells (Le Scolan *et al*., 2008). Finally, JUNB is an important regulator of TGFβ-induced invasive behavior and expression of EMT-related target genes (Sundqvist *et al*., 2018). Interestingly, the regulating TFs identified in this work globally perturb the regulation of most if not all TGFβ-dependent target genes, thereby exhibiting limited specificity in controlling the gene expression program. For instance, JUNB as an activator globally boosts both the up- and downregulated genes, thereby acting as an amplifier of TGFβ-dependent gene regulation irrespective of the direction of change. On the contrary, the repressors dampen the TGFβ response, simultaneously reducing the vast majority of up- and downregulated genes. Of note, some TFs, i.e., JUNB, SNAI1, SNAI2 or RUNX1, do show signs of specific regulation, as their KD affects subsets of target genes much more than the rest (Supplementary Figure 6, gray dots). Hence, these are strong candidates for controlling specific cell fates, e.g., by predominantly controlling EMT genes, while having lesser effects on other targets. Notably, in terms of TGFβ dose discrimination, specificity can also arise by TFs homogeneously affecting all target genes at specific time points. For instance, the observed early repression and late activation of target genes by the tested TFs may ensure that a common global gene expression program is dampened for a transient SMAD input (low TGFβ dose) but amplified for a sustained SMAD signal (high TGFβ dose), thereby ensuring specific gene expression response at the high dose.

Complex target gene expression kinetics e.g., delayed, continuous, biphasic, immediate early and inconsistent kinetics could not be fitted by our SMAD input model, and therefore require regulation by other dynamically changing TFs, as implemented in our extended FFL models. In similarity to previous work on EGF-induced transcriptional networks (Amit *et al*., 2007), we provide evidence for feedforward regulation in TGFβ-dependent gene expression, as late target gene expression requires ongoing SMAD signaling (Figure 4, C, Supplementary Figure, 3, D, E, F) and is particularly sensitive to the KD of induced TFs cooperating with SMAD proteins. In line with feedforward regulation by JUNB, it has been reported that JUNB redirects SMAD2 binding to new target gene promoters 16 h post-stimulation (Sundqvist *et al*., 2018), and that late target genes (6 h) are enriched for both SMAD and AP-1 TF binding motifs (Antón-García *et al*., 2023). Interestingly, in our data, both amplifying and repressing FFLs seem to mainly operate upon high dose stimulation, as twice as many TFs were induced by a strong stimulus compared to low dose (Supplementary Figure, 3, A). This suggests that FFLs mediate the late amplification of DDGs and selective EMT induction for strong TGFβ stimulation.

In summary, we employed a systems biology approach, combining ODE modeling with time-resolved RNA sequencing and phenotypic assays to obtain a holistic understanding of how feedforward regulation and SMAD signaling dynamics shape target gene expression. We demonstrate that the temporal dynamics of the SMAD signal and co-regulators of SMAD proteins shape the invasive behavior of cells. Therefore, our work points to potential intervention targets in malignancy and may serve as a blueprint for future quantitative studies linking signaling pathway dynamics to specific gene expression responses and cell fate decisions.

## Materials and Methods

### Cell culture

MCF10A cells were cultured in DMEM/F12 medium (Thermo Fisher, #21331020) supplemented with 100 ng/ml cholera toxin (Sigma, #C8052), 20 ng/ml epidermal growth factor (EGF) (Preprotech, #AF-100-15), 10 ug/ml insulin (Sigma, #I9278), 500 ng/ml hydrocortisone (Sigma, #H0888), GlutaMax (Thermo Fisher, #35050038), 5 % horse serum (Thermo Fisher, #16050122), and 1x penicillin/streptomycin (Thermo Fisher, #15140122). To culture H2B and SMAD2-tagged MCF10A cells, 200 ug/ml Geneticin (Thermo Fisher, #10131035) and 20-50 ug/ml Hygromycin (Thermo Fisher, #10687010) was added to the medium. Cells were trypsinized for ∼15 min with 0.05 % Trypsin/EDTA (Thermo Fisher, #15400054).

### Ligand and inhibitor treatment

Lyophilized TGFβ-1 derived from CHO-cells (Prepotech, # 100-21C) was reconstituted in 0.1 % BSA and 4 mM HCl. MCF10A cells were cultured 2 d before stimulation with different doses of TGFβ. To stop TGFβ-signaling a TGFβ-receptor-inhibitor (SB431542) (Merck, #616461) blocking the kinase activity of TGFβ –receptor-II was applied 90 min or 300 min post-stimulation with a final concentration of 10 μM (Figure 4, Supplementary Figure, 3). In long-term stimulation experiments, old medium was aspirated and cells were re-stimulated every 48 h with TGFβ-1.

### SMAD2 nuclear translocation data

SMAD2 nuclear translocation data for 351 and 378 single cells upon 2.5 and 100 pM TGFβ stimulation, respectively, with a sampling rate of every 5 min over a period of 24 h, was originally published in Strasen *et al*., 2018 (Strasen *et al*., 2018) (Figure 1). Here, we only considered the population-median at each time point across all individual cells, therefore neglecting cellular heterogeneity. Furthermore, the original data, given as a nuclear-to-cytoplasmic ratio of the SMAD2 protein, was background-subtracted to provide a zero-baseline input for mathematical modeling (Figure 3).

### siRNA mediated knock-down

siRNA mediated KD was performed using Dharmacon ON-TARGETplus Human siRNA SMART pools containing a selection of four different siRNA sequences targeting the gene of interest (*SMAD2: #L-003561-00-0005, SMAD3: #L-020067-00-0005, SNAI1: #L-010847-00-0005, SNAI2: #L-017386-00-0005, RUNX1: #L-003926-00-0005, SKIL: # L-010535-00-0005, SKI: #L-003927-00-0005, JUNB: #L-003269-00-0005, JUN: #L-003268-00-0005, KLF10: #L-006566-00-0005, ATF3: #L-008663-00-0005*). A pool of non-targeting siRNAs was used as negative control (Dharmacon, #D-001810-10). Reverse transfection with RNAiMAX (Lipofectamine™ RNAiMAX, #13778075) was performed at 10 nM final siRNA concentration. For 2x10^6^ cells in 10 ml cell culture medium lacking any antibiotics, 500 ul Opti-MEM™ (#1985062), 5 ul siRNA (stock concentration: 20 uM) and 10 ul RNAiMAX were pre-incubated for 20 min at RT before transfection. For TGFβ stimulation experiment, MCF10A cells were stimulated with 100 pM TGFβ 24 h post-KD generation and harvested at the time of stimulation (0 h) and later time points (90/ 180/ 360/ 720 min post-stimulation) (Figure 5, Supplementary Figure 5, 6). To conduct phenotypic analysis on KDs (see Flow cytometry analysis), the siRNA KD was renewed every 48 h (Figure 5, E, Supplementary Figure 7).

### bulk RNA sequencing and qPCR

RNA was isolated using the RNeasy Plus RNA-isolation kit (Qiagen, #74136) according manufactures instructions. Quantification of RNA concentrations and purity were determined with the Nanodrop. RNA quality was assessed and analyzed using RNA ScreenTapes and specific sample buffer (#RNA-Screen Tapes #5067-5576, RNA Screen Tape sample buffer #5067-5577, RNA Screen-Tape Ladder #5067-5578) in the Agilent TapeStation device. Calculated RINe values > 6 with flat baseline were strived for RNA sequencing experiments. Library preparation and bulk RNA sequencing experiments were performed in biological triplicates by Novogene (Figure 4, 5, 6) using poly-A-enrichment and paired-end-sequencing of 150 bp a NovaSeq6000 instrument with collecting 20 mio reads.

Library preparation of time course bulk RNA sequencing experiments upon low and high dose of TGFβ and initial data processing were performed in biological triplicates by the IMB Genomics Core Facility (Mainz) (Figure 1, 2). Following poly-A-enrichment transcripts were sequenced by single-end-sequencing of 75 bp on the NextSeq 500 sequencing platform.

Quantitative PCR (qPCR) was performed in a two-step process. First, the SuperScript™ Reverse Transcriptase kit was used to generate cDNA according the manufactures instruction (Thermo Fisher, #18064014) using Olig(dT)-primer for mRNA specificity. Specific forward and reverse primers for target genes of interest were designed and purchased from IDT (SMAD2; FWD: 5’ GGGTTTTGAAGCCGTCTATCAGC 3’, REV: 5’ CCAACCACTGTAGAGGTCCATTC 3’, SMAD3; FWD: 5’ CATCGAGCCCCAGAGCAATA 3’, REV: 5’ GTAACTGGCTGCAGGTCCAA 3’, SNAI1; FWD: 5’ GCCTAGCGAGTGGTTCTTCT 3’, REV: 5’ TGCTGGAAGGTAAACTCTGGATT 3’, SNAI2; FWD: 5’ TTCAACGCCTCCAAAAAGCC 3’, REV: 5’ AGAGATACGGGGAAATAATCACTGT 3’, RUNX1; FWD: 5’ CGTGGTCCTACGATCAGTCC 3’, REV: 5’ GTCGGGTGCCGTTGAGA 3’, SKIL; FWD: 5’ GGCTGAATATGCAGGACAG 3’, REV: 5’ TGAGTTCATCTTGGAGTTCTTG 3’, SKI; FWD: 5’ TTCCGAAAAGGACAAGCCGT 3’, REV: 5’ ACAGCCCAGGCTCTTATTGG 3’, JUNB; FWD: 5’ CCTGGACGATCTGCACAAGA 3’, REV: 5’ GTAGCTGCTGAGGTTGGTGT 3’, JUN; FWD: 5’ CCTGGACGATCTGCACAAGA 3’, REV: 5’ GTAGCTGCTGAGGTTGGTGT 3’, KLF10; FWD: 5’ CGAGGACGCACACAGGAGAA 3’, REV: 5’ GTCACTCCTCATGAACCGCC 3’, ATF3; FWD: 5’ TCCATCACAAAAGCCGAGGTAG 3’, REV: 5’ CTGCAGGCACTCCGTCTTC 3’, SERPINE1: FWD: 5’ GGCTGACTTCACGAGTCTTTCA 3’, REV: 5’ ATGCGGGCTGAGACTATGACA 3’, FN1: FWD: 5’ GTGTGATCCCGTCGACCAAT 3’, REV: 5’ CGACAGGACCACTTGAGCTT 3’, SMAD7: FWD: 5’ ACCCGATGGATTTTCTCAAACC 3’, REV: 5’ GCC AGA TAA TTCGTT CCC CCT 3’, AXL: FWD: 5’ ACTCTGGGAGAGGGAGAGTT 3’, REV: 5’ GAGCCTCATGACGTTGGGAT 3’) and used at final concentrations of 240 nM. Second, for qPCR the Power SYBR ® Green PCR Master Mix (Thermo Fisher, #4367659) was used at 1x final concentration having SYBR ® Green I Dye, AmpliTaq Gold ® DNA Polymerase, dNTPs and internal passive reference dye (ROX), included. The qPCR was performed on a QuantStudio™ 5 Real-Time-PCR-System (Figure 6, C, Supplementary Figure 5, A; 7, A).

### Flow cytometry analysis

The expression levels of Vimentin and E-Cadherin were assessed at the protein level by flow-cytometry (Figure 2). MCF10A cells were treated for 5 or 8 d with TGF-β1 (1/ 5/ 50/ 250 pM or 100 pM), with subsequent restimulation every 48 h. To assess the impact of co-factors on EMT initiation, cells were first transfected and then stimulated 24 h later for 8 d with 100 pM TGF-β1 with subsequent restimulation every 48 h (Figure 5, Supplementary Figure 7). Cells were harvested by TrypLE™ (ThermoFisher, #12604013). Before fixation, cells were stained with LIVE/DEAD™ Fixable Dead Cell Stain for 30 min at 4 °C (Thermo Fisher, #L34975) to differentiate between living and dead cells. For cell fixation and permeabilization, cells were treated with 4 % PFA in DPBS for 15 min, and 0.1 % Triton in FACS media (DPBS + 5 % FHS) for 15 min at 4 °C. Cells were then incubated with anti-CD324 (E-Cadherin) antibody (Cell signaling, #14472, clone 4A2, monoclonal mouse Ab, 1:150) and anti-Vimentin antibody (Cell signaling, #5741, clone D21H3, monoclonal rabbit Ab, 1:150) for 1 h. After blocking in FACS media for 10 min, cells were incubated with secondary antibodies Alexa Fluor 647 goat anti-mouse IgG (H+L), (Thermo Fisher, #2369432, 1:1000) and Alexa Fluor 488 goat anti-rabbit IgG (H+L) (Thermo Fisher, #2382186, 1:250) for 45 min. All antibody incubations were performed in FACS media. Cells were washed between live-dead-staining and fixation with DPBS and during antibody staining with FACS media for two times and analyzed by MACSQuant10 flow-cytometer. Flow cytometry data was analyzed by FlowJo v10.8.1, quantifying FITC-A SSC-A (Vimentin) or APC-A SSC-A (E-Cadherin) in the cell population.

### Bioinformatics analysis of bulk RNA sequencing data

Basic bioinformatics analysis from raw sequencing files to DESeq2 was done by the bioinformatics core facility at the IMB Mainz (*time course data*) and Novogene (*KD data*). Raw sequence reads were assessed with FastQC (v. 0.11.5) (S., 2010) and aligned to the human reference genome GRCh38.p12 (GTF annotation file from Gencode human release 25) using STAT aligner (v. 2.5.2b) or Hisat2 aligner (v. 2.0.5) (Dobin *et al*., 2013) (Kim, Langmead and Salzberg, 2015) for the KD data, respectively. Mapped data were summarized on the gene level using the featureCounts software (v.1.5.0-p3) (Liao, Smyth and Shi, 2014). Further data evaluation, normalization and pairwise differential expression analyses were carried out in R using the Bioconductor package DESeq2 (*v*.*1*.*18*.*1, time course data, v*.*1*.*20*.*0, KD data*) (Love, Huber and Anders, 2014). To investigate the number of genes, sign. diff. regulated transcripts were filtered for protein coding genes and RPKM/ FPKM > 0.5. Differentially expressed genes were isolated based on FDR < 0.01 and absolute fold change ≥ 1.5.

Phenotypic specific gene groups were selected from different sources (*see Supplementary table 1 and 3*); EMT genes: curated general EMT gene list (Zhao *et al*., 2015) and the TGFβ-EMT gene signature (Gordian *et al*., 2019), cell cycle genes: https://www.gsea-msigdb.org/gsea/msigdb/cards/KEGG_CELL_CYCLE

### Implementation of a minimal gene expression model

Ordinary differential equation (ODE) models were used to describe the effect of SMAD2 on the dynamics of target gene expression. As an input for the model, we used background-subtracted microscopy data (see SMAD2 nuclear translocation data) with spline interpolation between 5 min sampling intervals. Transcriptional regulation by SMAD was modeled using Hill function and two possible cases were considered-activation and inhibition of target genes by SMAD. The ODE for the activation model reads

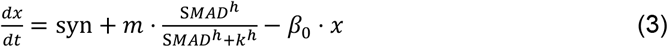

where *x* is the concentration of target gene mRNA, whereas syn and *β*_0_ are the basal synthesis and degradation rates, respectively. In the Hill term, SMAD is the nuclear concentration of SMAD, *m* = is the maximal SMAD induced transcriptional rate, *h* the Hill coefficient, and *k* the half-saturation point. To directly compare our model to RNA sequencing data, which quantifies the fold change rather than concentration of mRNA in response to TGFβ stimulation, we divided both sides of Equation 3 by the steady-state mRNA level without TGFβ stimulation, and the steady-state solution can be derived by the following

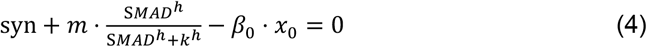

without TGFβ stimulation, SMAD=0, so we have,

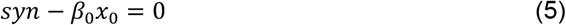

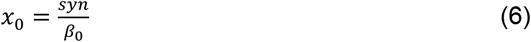

We then divided both sides of Equation 3 by *x*_0_:

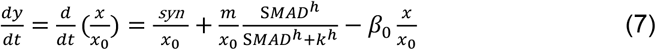

where the steady-state expression 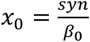 can be used to substitute 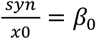 and 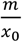 replaced with *k*_*syn*_

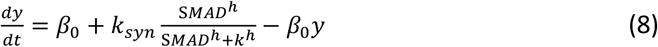

We can then use the above ODE solution to directly compare with the time-resolved RNA sequencing data. The same principle was used for the inhibition model (see supplementary methods). We fit both activation and inhibition models independently to each gene by minimizing the chi-square (*χ*^2^) (Sheskin, 2003; Kanji, 2006) value defined as,

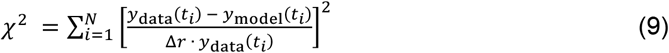

where *y*_data_(*t*_*i*_) and *y*_model_(*t*_*i*_) are the measured and modeled mRNA fold changes at time point *t*_*i*_, respectively. The RNA sequencing data were collected from 12 time points in total (6 for high dose and 6 for low dose of TGFβ) for each gene, i.e., *N* = 12. The open-source Matlab package Data2Dynamics (D2D) was used for model fitting (Raue *et al*., 2013, 2015). To avoid local minima, the Latin hypercube sampling (n = 500) was used to sufficiently scan the initial parameter space. To determine the relative error, we calculate the slope of a scatter plot, plotting mean RPKMs (a) vs. standard deviation (b) (see Supplementary Figure 4). We make use of this relative error (r) and use the Gaussian error propagation (see supplementary methods).

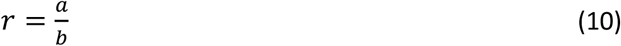

Using the calculated slope of the linear error model on RPKMs (m= 0.082) (see Supplementary Figure 4), the Gaussian error propagation equation results in a relative error of 0.11; 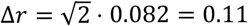

For both the activation and inhibition models, we employed a *χ*^2^ goodness-of-fit test (Cedersund and Roll, 2009) to assess their agreement with the experimental data. The decision to accept or reject a model was determined by comparing the *χ*^2^ value from the best fit to the critical *χ*^2^ value given by the *χ*^2^ distribution (with corresponding degrees of freedom) at 95 % confidence level. The degrees of freedom are 8 (12 data points minus 4 free parameters) for the activation model and 9 (12 data points minus 3 free parameters) for the inhibition model. The critical *χ*^2^ values were established as 15.5 for the activation model and 16.9 for the inhibition model. If the fitted *χ*^2^ is lower than the corresponding critical *χ*^2^ value, the model is accepted; otherwise, the model is rejected. In case that both activation and inhibition models were accepted by the *χ*^2^ test for a gene, we used the Akaike information criteria (AIC) (Akaike, 1974) for further model discrimination,

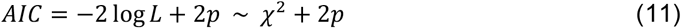

where *L* is the likelihood and *p* the number of free parameters. Because here we assume normally distributed errors, the *χ*^2^ value and −2log*L* only differ by a constant value and therefore can be used to calculate AIC for different models. The model with the lower AIC was selected. Based on these criteria, we were able to describe 19% of target genes by the activation model and 45% by the inhibition model. Both models were rejected for the remaining 36% (1744 out of 4823) target genes (Figure 3).

### Model extension by feed-forward loops

To explain the genes rejected by the simple activation and inhibition models (see above), we implemented extended models taking into account feed-forward loops (FFLs) (Mangan and Alon, 2003). We introduced an unknown regulator, named TF, which is activated directly by SMAD. TF regulates the target gene (TG), as either an activator or a repressor, jointly with SMAD via AND or OR gate-logic. This generates in total 8 different models (Figure 4, D, Supplementary Figure 3, G). There are two equations for each of the 8 model variants. The first equation describes the activation of TF and is similar to the simple activation model (1).

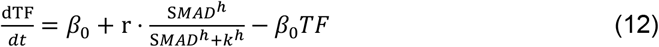

The second equation differs among the 8 models depending on activating or repressor role of TF and SMAD. In addition, the transcription rate changes if we assume an AND gate = f(SMAD) x f(TF) or OR gate-logic f(SMAD) + f(TF) − f(SMAD) x f(TF) (Rogers *et al*., 2022) (see supplementary methods for initial FFL equations and their steady-state). By dividing the equations by the steady-state (Supplementary Methods), we obtain the following equations for each model:

Model 1: AND-gate: SMAD activator, TF activator

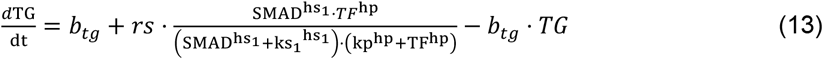

Model 2: OR-gate: SMAD activator, TF activator

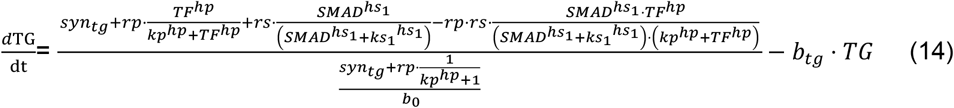

Model 3: AND-gate: SMAD repressor, TF activator

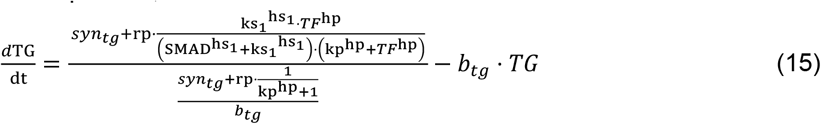

Model 4: OR-gate: SMAD repressor, TF activator

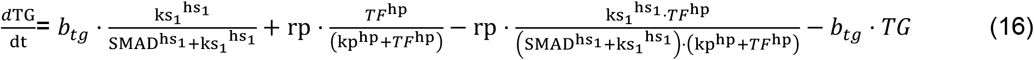

Model 5: AND-gate: SMAD activator, TF repressor

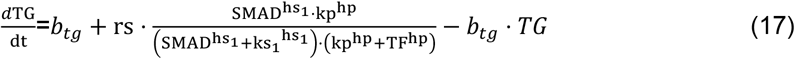

Model 6: OR-gate: SMAD activator, TF repressor

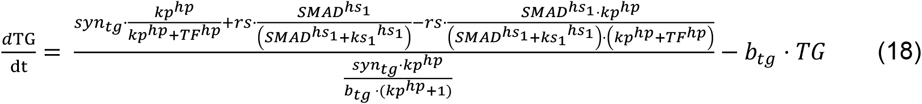

Model 7: AND-gate: SMAD repressor, TF repressor

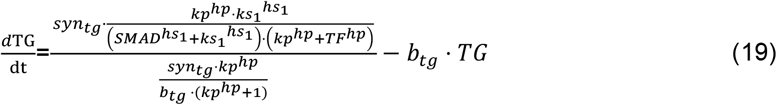

Model 8: OR-gate: SMAD repressor, TF repressor

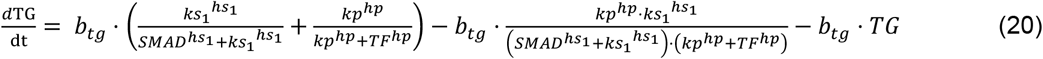

where the parameters are defined as follows:

*syn*_*tg*_ = basal target gene synthesis rate

*rs* = maximal rate of SMAD-induced target gene (TG) transcription

rp = maximal rate of TF-induced target gene transcription

*hs*_1_= hill coefficient of SMAD-induced target gene transcription

ks_1_= half-saturation point of SMAD regulation

*hp* = hill coefficient of TF-induced target gene transcription

*kp* = half-saturation point of TF regulation

*b*_*tg*_= degradation rate of target gene mRNA

The 8 FFL models were fitted to individual rejected genes using the same method as applied earlier for the simple model. As shown above, the model equations 13-20 for the FFL possess many parameters (up to 12). To obtain the effective degrees of freedom (DOF) for the *χ*^2^ test, we applied a simulation-based method proposed by Kreutz (Kreutz, 2020) (see Supplementary Figure 4B). To determine the DOF for each of the 8 FFL model, we generated synthetic data from the best fit and gain a model-intrinsic *χ*^2^ by refitting.

First, we selected 14 rejected genes (by simple models) that represent different dynamic groups (Figure 4). Then we fit 8 FFL models to each of them and selected the best model using AIC. For each gene, we added normally distributed noise with 11 % standard deviation to the time trajectories of the best fit, generating 100 synthetic data sets (Supplementary Figure 4, B). We then refit the model to the 100 samples and calculated the *χ*^2^. The *χ*^2^ values before refit is quantified as:

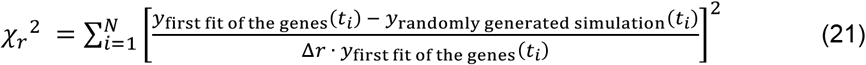

and after refit:

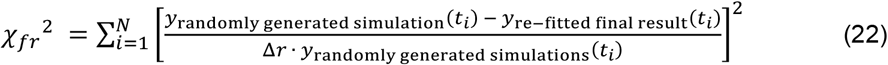

We calculated the model-intrinsic degrees of freedom following the formula *DOF* = *χ*_*r*_^2^ − *χ*_*fr*_^2^ = 12 − 6 = 6 (Supplementary Figure 4, C). Based on this effective degrees of freedom, we performed model rejection/selection on all 1744 genes (rejected by the simple models) using *χ*^2^ test or/and AIC.

### Classification of rejected genes

To investigate why the simple model is not able to explain 1744/ 4823 differentially expressed genes, we analyzed the gene expression kinetics shared by the rejected genes and are rarely found in the described genes. For that, we have grouped rejected genes into 5 subgroups (Figure 4) “delayed expression kinetics”, “inconsistent expression kinetics”, “biphasic expression kinetics”, “immediate early expression kinetics” and “continuous expression kinetics”, by using different filter settings as indicated in Supplementary Table 4.

### Identification of candidate TFs mediating feedforward regulation

We have used an earlier published (https://www.sciencedirect.com/science/article/pii/S0092867418301065) list of described TFs (Lambert *et al*., 2018). Across all early time points (<360 min) 99 TFs are differentially expressed upon low dose stimulation and 133 TFs are differentially expressed upon high dose stimulation. We further isolated TFs, which are following the SMAD dynamics and are therefore SMAD induced TF and share differentially expression between 45-90 min post-stimulation. To further narrow the list of TFs, we filtered co-factors already described in literature to interact with SMADs. This result in a list of 6 co-factors (SNAI2, JUN, JUNB, SMAD7, KLF10, SKIL) differentially expressed upon low dose application (Figure 5). The same 6 co-factors and 4 additional ones (RUNX1, SKI, E2F5, KDM6B) are differentially expressed upon high dose application (Figure 5). We have chosen only the upregulated TFs to focus on (SNAI2, RUNX1, SKIL, SKI, JUNB, JUN, KLF10) and added ATF3, SNAI1 as well as SMAD2 and SMAD3 as positive controls.

### Assessing impact of co-factor KD on target gene expression (slope quantification)

To divide SMAD co-factors into activators and repressors, we quantified the slope of scatter plots with gene expression values of the non-targeting control (NTC) plotted on the x-axis against gene expression values of the KD (KD) condition shown on the y-axis (Figure 5, Supplementary Figure 6). With this analysis we quantified whether the gene expression is on average, stronger or weaker upon KD compared to control. For further analysis, we only selected KDs with significantly changed slopes ≠ 1 (t-test, α=0.05). Furthermore, we identified the variance among the three biological replicates by plotting their FPKM-values in scatter plots against each other (Supplementary Table 6).

### Inference of transcription rate upon JUNB KD

To infer the time-dependent transcription rate without prior knowledge of molecular interactions among SMAD and cofactors, such as JUNB, we started with a generic ordinary differential equation describing the kinetics of fold change in mRNA expression (*x*),

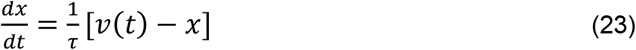

where *v*(*t*) is the TGFβ-stimulated transcription rate relative to the basal transcription rate and thus *v*(0) = 1. *t* is the mRNA lifetime. By rearranging the equation 23, we obtained the transcription rate *v*(*t*),

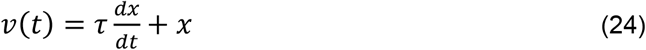

Equation 24 suggests that the transcription rate *v*(*t*) can be directly inferred from the time-resolved RNA sequencing data. To overcome that the calculation of the time derivative (d*x*/d*t*) is hindered by sparse time points, we used piecewise cubic-spline interpolation to yield interpolated mRNA trajectories with higher time resolution using a 15 min sampling interval. To avoid over flexibility of the naïve cubic splines, we used the modified Akima interpolation method implemented in Matlab (makima). We then calculated numerically the discrete time derivative (Δ*x*/Δ*t*) for each interpolated time point. For the value of mRNA lifetime τ, we used a fixed value of 9 h for all mRNAs, corresponding the median mRNA lifetime in mammalian cells (Schwanhäusser *et al*., 2011). Notably, according to the Equation 24, the lifetime is just a scaling factor of the time derivative, mainly controlling the amplitude of the inferred transcription rate *v*(*t*) with little effects on its temporal shape. To quantitatively understand the regulatory role of JUNB, we separately performed the transcription-rate inference for each gene belonging to the DDG, delayed and biphasic gene groups and calculated the difference of transcription rates between JUNB KD and control conditions (Δ*v*(*t*) = *v*_KD_ − *v*_control_). For each group, we then calculated the median transcription rate difference Δ*v*(*t*) at each time point, separately considering the upregulated (Figure 6, C) and downregulated (Supplementary Figure 8, C) genes (defined based on their direction of change at the final time point (720 min) in the control condition).

## Acknowledgments

We thank Monilola Olayioye and Merih Özverin (Institute of Cell Biology and Immunology, University of Stuttgart) for the help with flow cytometry analysis, the core facility at the IMB in Mainz for performing RNA sequencing and Clemens Kreutz for the discussion on model acceptance analysis. The authors acknowledge the support by the state of Baden-Württemberg through bwHPC. Portions of this work were developed from the doctoral dissertations of L.Hartmann and P.K. This work was supported by DFG funding to A.L. (LO 1634/7-1) and S.L. (LE 3473/4-1).

