## Supplementary Information for "Transcriptional regulators ensuring specific gene expression and decision making at high TGFβ doses"

**This PDF file includes:**

Supporting text
Figures S1 to S8
Tables S1 to S7
SI References

**Supporting Information Text**

**Methods**

**Implementation of a minimal gene expression model: Inhibition model**

The same principle used for the activation model (see methods) was used for the inhibition model. The equation for the inhibition model is:

$$\frac{dx}{dt} = \mathrm{syn} \cdot \frac{k^h}{SMAD^h + k^h} - \beta_0 \cdot x \quad \text{(S1)}$$

When t=0 and SMAD=0 and therefore $\frac{k^h}{SMAD^h+k^h} = 1$, the steady state is defined by:

$$0 = \mathrm{syn} \cdot 1 - \beta_0 \cdot x0 \quad \text{(S2)}$$

$$x_0 = \frac{syn}{\beta_0} \quad \text{(S3)}$$

We define the mRNA expression fold change $y \equiv \frac{x}{x_0}$. The respective ODE can be derived from S1:

$$\frac{dy}{dt} = \frac{d}{dt}\left(\frac{x}{x_0}\right) = \frac{syn}{x_0} \cdot \frac{k^h}{SMAD^h+k^h} - \beta_0 \cdot \frac{x}{x_0} \quad \text{(S4)}$$

where the steady-state expression $x_0 = \frac{syn}{\beta_0}$ can be used to substitute $\beta_0 = \frac{syn}{x_0}$

$$\frac{dy}{dt} = \beta_0 \cdot \frac{k^h}{SMAD^h+k^h} - \beta_0 \cdot y \quad \text{(S5)}$$

Or equivalently,

$$\frac{dy}{dt} = \beta_0 - \beta_0 \cdot \frac{SMAD^h}{SMAD^h+k^h} - \beta_0 \cdot y \quad \text{(S6)}$$

To calculate the error, we use the Gaussian Error propagation:

$$\Delta F = \sqrt{\sum_x \left(\frac{\partial F}{\partial x}\right)^2 \Delta x^2} \quad \text{(S7)}$$

Where F is the fold change, r=a/b, and x=(a,b)

$$\Delta r = \sqrt{\left(\frac{\partial r}{\partial a}\right)^2 \cdot \Delta a^2 + \left(\frac{\partial r}{\partial b}\right)^2 \cdot \Delta b}$$
$$\frac{\partial r}{\partial a} = \frac{1}{b}; \frac{\partial r}{\partial b} = -\frac{a}{b^2} \quad \text{(S8)}$$

$$\Delta r = \sqrt{\frac{\Delta a^2}{b^2} + \frac{a^2 \cdot \Delta b^2}{b^4}} = \sqrt{\frac{\Delta a^2 \cdot b^2}{b^4} + \frac{a^2 \Delta b^2}{b^4}} \quad \text{(S9)}$$

Using the slope m of the linear error model on RPKMs which is defined by the absolute values a and b instead of the relative r, we substitute $\Delta a = m \cdot \mathrm{a}$ ; and $\Delta b = m \cdot b$ and receive

$$\Delta r = \sqrt{\frac{m^2 \cdot a^2 \cdot b^2}{b^4} + \frac{a^2 \cdot m^2 \cdot b^2}{b^4}} = \sqrt{2 \cdot m^2 \cdot \left(\frac{a}{b}\right)^2} \quad \text{(S10)}$$

The following equation is used to calculate the relative error based on the calculated slope.

$$\Delta r = \sqrt{2 \cdot m^2 \cdot r^2} \quad \text{(S11)}$$

**Model extension by feed-forward loops:**

The equations for the 8 extended models considering FFL regulation in an AND-or OR-gate logic (SF 3, G) are listed below without normalization by steady-state:

Model 1: AND-gate: SMAD activator, TF activator

$$\frac{d\mathrm{TG}}{\mathrm{dt}} = syn_{tg} + rs \cdot \frac{\mathrm{SMAD}^{\mathrm{hs}_1} \cdot TF^{\mathrm{hp}}}{\left(\mathrm{SMAD}^{\mathrm{hs}_1} + {\mathrm{ks}_1}^{\mathrm{hs}_1}\right) \cdot \left(\mathrm{kp}^{\mathrm{hp}} + TF^{\mathrm{hp}}\right)} - b_{tg} \cdot TG \quad \text{(S12)}$$

Model 2: OR-gate: SMAD activator, TF activator

$$\frac{d\mathrm{TG}}{\mathrm{dt}} = syn_{tg} + \mathrm{rp} \cdot \frac{TF^{\mathrm{hp}}}{\mathrm{kp}^{\mathrm{hp}} + TF^{\mathrm{hp}}} + \mathrm{rs} \cdot \frac{\mathrm{SMAD}^{\mathrm{hs}_1}}{\left(\mathrm{SMAD}^{\mathrm{hs}_1} + {\mathrm{ks}_1}^{\mathrm{hs}_1}\right)} - \mathrm{rp} \cdot rs \cdot \frac{\mathrm{SMAD}^{\mathrm{hs}_1} \cdot TF^{\mathrm{hp}}}{\left(\mathrm{SMAD}^{\mathrm{hs}_1} + {\mathrm{ks}_1}^{\mathrm{hs}_1}\right) \cdot \left(\mathrm{kp}^{\mathrm{hp}} + TF^{\mathrm{hp}}\right)} - b_{tg} \cdot TG \quad \text{(S13)}$$

Model 3: AND-gate: SMAD repressor, TF activator

$$\frac{d\mathrm{TG}}{\mathrm{dt}} = syn_{tg} + \mathrm{rp} \cdot \frac{{\mathrm{ks}_1}^{\mathrm{hs}_1} \cdot TF^{\mathrm{hp}}}{\left(\mathrm{SMAD}^{\mathrm{hs}_1} + {\mathrm{ks}_1}^{\mathrm{hs}_1}\right) \cdot \left(\mathrm{kp}^{\mathrm{hp}} + \mathrm{TF}^{\mathrm{hp}}\right)} - b_{tg} \cdot TG \quad \text{(S14)}$$

Model 4: OR-gate: SMAD repressor, TF activator

$$\frac{d\mathrm{TG}}{\mathrm{dt}} = syn_{tg} \cdot \frac{{\mathrm{ks}_1}^{\mathrm{hs}_1}}{\mathrm{SMAD}^{\mathrm{hs}_1} + {\mathrm{ks}_1}^{\mathrm{hs}_1}} + \mathrm{rp} \cdot \frac{TF^{\mathrm{hp}}}{\left(\mathrm{kp}^{\mathrm{hp}} + TF^{\mathrm{hp}}\right)} - \mathrm{rp} \cdot \frac{{\mathrm{ks}_1}^{\mathrm{hs}_1} \cdot TF^{\mathrm{hp}}}{\left(\mathrm{SMAD}^{\mathrm{hs}_1} + {\mathrm{ks}_1}^{\mathrm{hs}_1}\right) \cdot \left(\mathrm{kp}^{\mathrm{hp}} + TF^{\mathrm{hp}}\right)} - b_{tg} \cdot TG \quad \text{(S15)}$$

*Model 5: AND-gate: SMAD activator, TF repressor*

$$\frac{d\mathrm{TG}}{\mathrm{dt}} = syn_{tg} + \mathrm{rs} \cdot \frac{\mathrm{SMAD}^{\mathrm{hs}_1} \cdot \mathrm{kp}^{\mathrm{hp}}}{\left(\mathrm{SMAD}^{\mathrm{hs}_1} + {\mathrm{ks}_1}^{\mathrm{hs}_1}\right) \cdot \left(\mathrm{kp}^{\mathrm{hp}} + TF^{\mathrm{hp}}\right)} - b_{tg} \cdot TG \quad \text{(S16)}$$

Model 6: OR-gate: SMAD activator, TF repressor

$$\frac{d\mathrm{TG}}{\mathrm{dt}} = syn_{tg} \cdot \frac{\mathrm{kp}^{\mathrm{hp}}}{\mathrm{kp}^{\mathrm{hp}} + TF^{\mathrm{hp}}} + \mathrm{rs} \cdot \frac{\mathrm{SMAD}^{\mathrm{hs}_1}}{\left(\mathrm{SMAD}^{\mathrm{hs}_1} + {\mathrm{ks}_1}^{\mathrm{hs}_1}\right)} - \mathrm{rs} \cdot \frac{\mathrm{SMAD}^{\mathrm{hs}_1} \cdot \mathrm{kp}^{\mathrm{hp}}}{\left(\mathrm{SMAD}^{\mathrm{hs}_1} + {\mathrm{ks}_1}^{\mathrm{hs}_1}\right) \cdot \left(\mathrm{kp}^{\mathrm{hp}} + TF^{\mathrm{hp}}\right)} - b_{tg} \cdot TG \quad \text{(S17)}$$

Model 7: AND-gate: SMAD repressor, TF repressor

$$\frac{d\mathrm{TG}}{\mathrm{dt}} = syn_{tg} \cdot \frac{\mathrm{kp}^{\mathrm{hp}} \cdot {\mathrm{ks}_1}^{\mathrm{hs}_1}}{\left(\mathrm{SMAD}^{\mathrm{hs}_1} + {\mathrm{ks}_1}^{\mathrm{hs}_1}\right) \cdot \left(\mathrm{kp}^{\mathrm{hp}} + \mathrm{TF}^{\mathrm{hp}}\right)} - b_{tg} \cdot TG \quad \text{(S18)}$$

Model 8: OR-gate: SMAD repressor, TF repressor

$$\frac{d\mathrm{TG}}{\mathrm{dt}} = syn_{tg} \cdot \left(\frac{{\mathrm{ks}_1}^{\mathrm{hs}_1}}{\mathrm{SMAD}^{\mathrm{hs}_1} + {\mathrm{ks}_1}^{\mathrm{hs}_1}} + \frac{\mathrm{kp}^{\mathrm{hp}}}{\mathrm{kp}^{\mathrm{hp}} + \mathrm{TF}^{\mathrm{hp}}}\right) - syn_{tg} \cdot \frac{\mathrm{kp}^{\mathrm{hp}} \cdot {\mathrm{ks}_1}^{\mathrm{hs}_1}}{\left(\mathrm{SMAD}^{\mathrm{hs}_1} + {\mathrm{ks}_1}^{\mathrm{hs}_1}\right) \cdot \left(\mathrm{kp}^{\mathrm{hp}} + \mathrm{TF}^{\mathrm{hp}}\right)} - b_{tg} \cdot TG \quad \text{(S19)}$$

Similar to the simple model, we calculated the steady-state without TGFβ stimulation. For t=0 and SMAD=0 we get the following steady-states per model:

Model 1: AND-gate: SMAD activator, TF activator

$$\frac{syn_{tg}}{b0} \tag{S20}$$

Model 2: OR-gate: SMAD activator, TF activator

$$\frac{syn_{tg}+\frac{rp}{kp^{hp}+1}}{b_{tg}} \tag{S21}$$

Model 3: AND-gate: SMAD repressor, TF activator

$$\frac{syn_{tg}+\frac{rp1}{kp^{hp}+1}}{b_{tg}} \tag{S22}$$

Model 4: OR-gate: SMAD repressor, TF activator

$$\frac{syn_{tg}}{b_{tg}} \tag{S23}$$

Model 5: AND-gate: SMAD activator, TF repressor

$$\frac{syn}{b_{tg}} \tag{S24}$$

Model 6: OR-gate: SMAD activator, TF repressor

$$\frac{syn_{tg}\cdot kp^{hp}}{b_{tg}\cdot(kp^{hp}+1)} \tag{S25}$$

Model 7: AND-gate: SMAD repressor, TF repressor

$$\frac{syn_{tg}\cdot kp^{hp}}{b_{tg}\cdot(kp^{hp}+1)} \tag{S26}$$

Model 8: OR-gate: SMAD repressor, TF repressor

$$\frac{syn_{tg}}{b_{tg}} \tag{S27}$$

The final equations used for FFL models (Figure 4, D, E. Supplementary table 5) are shown in methods: *Feed-Forward loops equation.*

**SMAD2 live-cell imaging upon TGFβ stimulated transcription factor KD cells**

For live-cell time-lapse microscopy, cells were plated and transfected with the corresponding siRNA, as described in methods. For this approach, 1x$10^5$ cells in 1 ml cell culture medium lacking any antibiotics, 50 ul Opti- MEMTM (#1985062), 0.5 ul siRNA (stock concentration: 20 uM) and 1 ul RNAiMAX, were seeded in a polymer coverslip bottom 24 well plate (ibidi) 2 d before the experiment. Before starting the experiment, cells were washed once with 1 × PBS, and the medium was changed to DMEM lacking phenol red, supplemented with all growth factors, 5 % horse serum, and antibiotics. To maintain constant $CO_2$ concentration (5 %), temperature (37 °C), and humidity, the microscope was placed in a custom incubator. Cells were imaged using a Nikon Ti inverted fluorescence microscope with a Nikon DS-Qi2 camera and a 20× plan apo objective (NA 0.75). The following filter sets were used: Venus (500/20 nm excitation, 515 nm dichroic beam splitter, 535/30 nm emission) and CFP (436/20 nm emission, 455 nm dichroic beam splitter, 480/40 nm excitation). Images were acquired every 10 min for the duration of the experiment using Nikon Elements software. TGFβ 1 was prepared in 125 µl media and added, after two loops of images, to achieve the final concentration in 625 µl media. See Supplementary Table 7 for measured cell numbers per condition.

### GSEA of DDGs

The g:Profiler was used to analyze enriched genes among the DDGs (Kolberg *et al.*, 2023). For multiple testing correction the build-in method g:SCS threshold was applied. We only considered KEGG and REACTOME gene sets and filtered for adjusted p-value < 0.01.

## S1

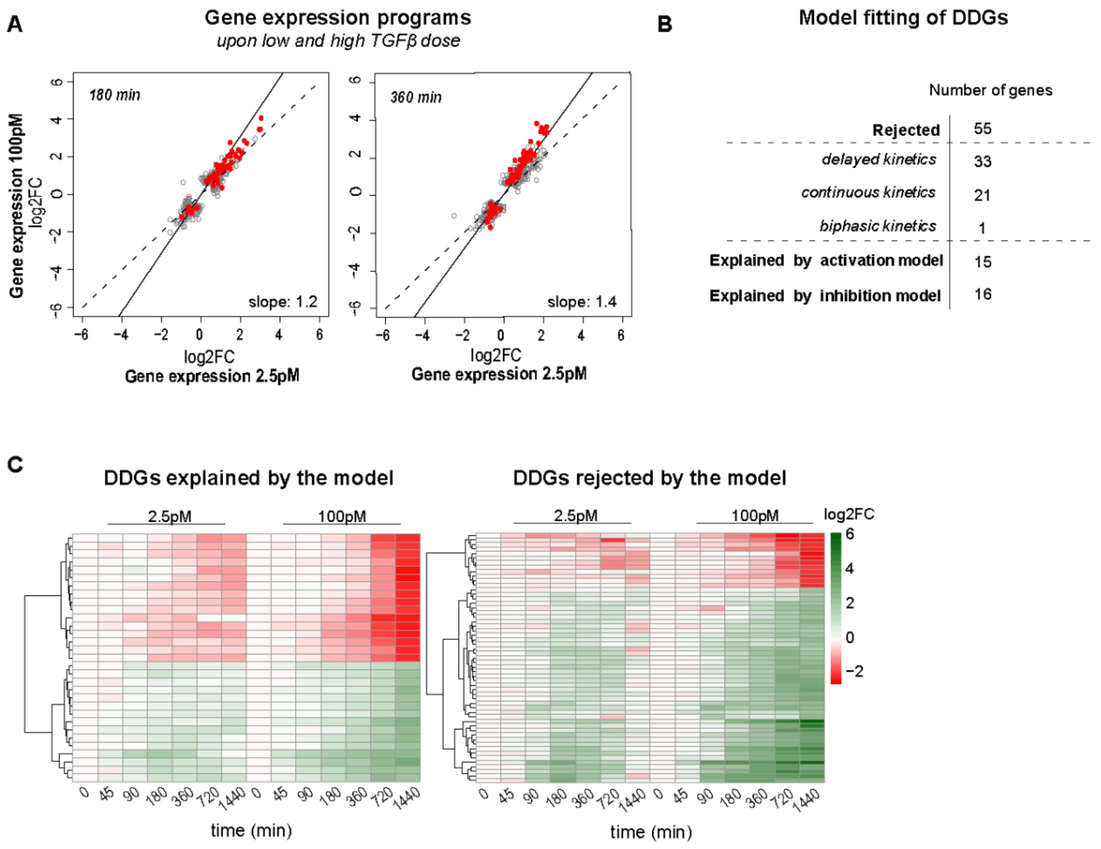


**S1: Features of dose-discriminating genes (DDGs),** *related to main Figure 2*

**A)** High and low doses of TGFβ induces similar gene expression programs early after stimulation. Scatter plot showing TGFβ-induced global gene expression changes upon low and high dose application at different time points (180/ 360 min) in MCF10A cells. Each grey dot represents a gene with significantly different expression (adj. p-value < 0.01, abs. FC > 1.5) compared to unstimulated cells in at least one of the two stimulation conditions. The solid line is a linear fit to the data, with the slope indicated in the bottom-right. Red dots indicate DDGs, which are differentially expressed upon high dose stimulation (absolute (abs.) fold-change (FC) >4*)* but not low dose stimulation (abs. FC<2) at late time points (720 or 1440 min post-stimulation). Number of genes shown in scatter plot: 180 min: DDGs: 58, all: 564, 360 min: DDGs: 73, all: 1069. **B)** The majority of DDGs (55/86) are rejected by the simple gene expression models (Fig. 3), whereas 15/86 and 16/86 are explained by the activation and inhibition models, respectively. The rejected DDGs belong to the gene sets with complex kinetics (Figure 4), i.e., the delayed (33 genes), continuous (21 genes) and biphasic (1 gene) groups. **C)** Heatmaps showing the expression dynamics of DDGs, separately for DDGs that can be explained by the simple gene expression models in Figure 3 (left) and those rejected by these models (right). Rejected genes show higher fold change values compared to explained ones, and are also characterized by a shutdown at the late time point (1440 min) at low dose stimulation.

**S2**

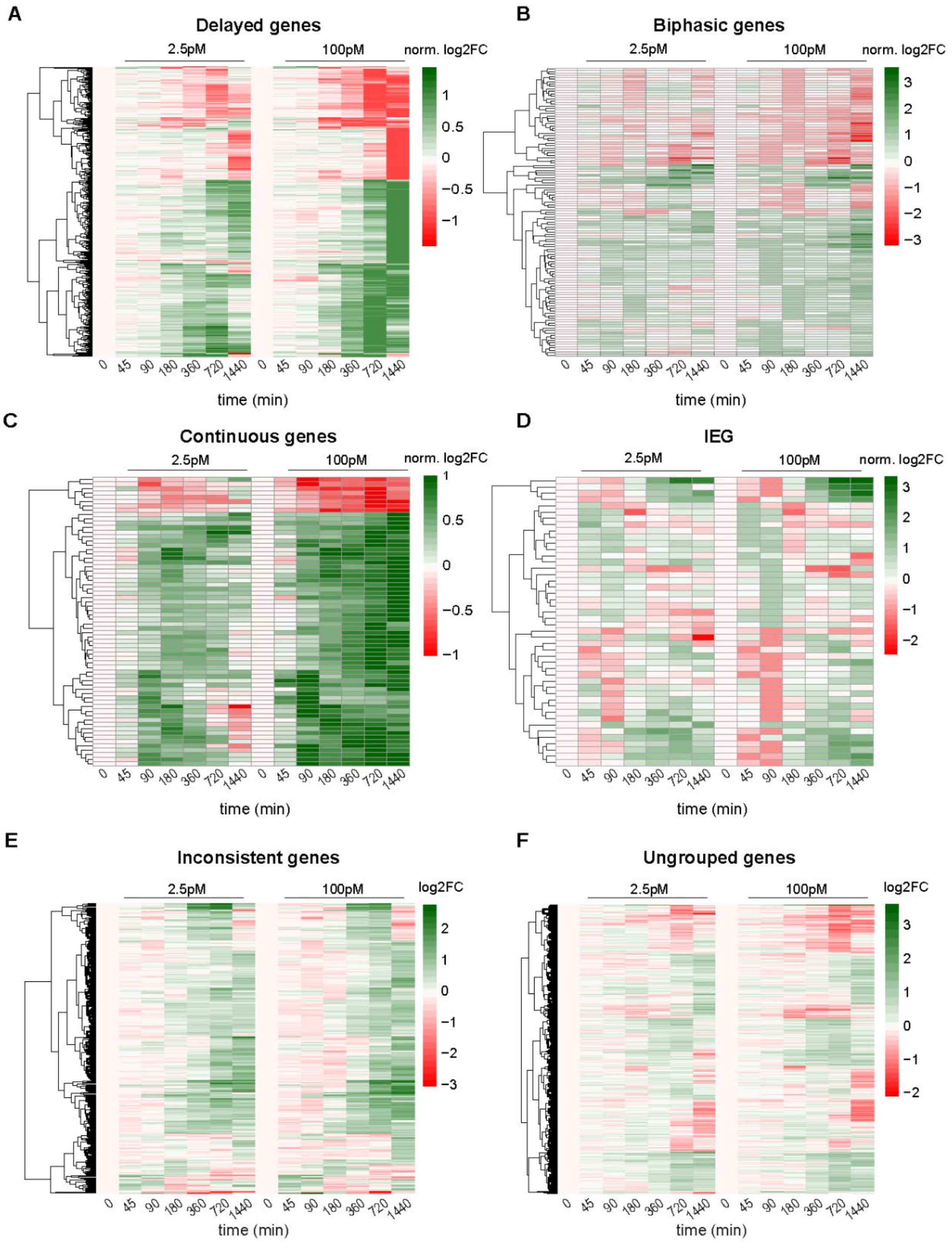


**S2: RNA sequencing time courses of target genes groups with complex kinetics,** *related to main Figure 4*

RNA sequencing time courses of all genes belonging to the gene groups in Figure 4. **A)** 483 delayed target genes. For better visibility of time course shapes, all log2 fold-changes of each time course were normalized by dividing by the highest log2-fold-change of that time course at one of the following time points (100 pM stimulation): 180/360/720 or 1440 min. Genes (y-axis) are sorted by hierarchical clustering**.** See Methods and Table S3 for filtering approach. **B)**112 biphasic target genes. Same as panel A, log2-fold-changes now being normalized to the highest absolute expression value after 45/ 90 or 180 min of 100 pM stimulation. **C)** 66 continuous target genes. Same as panel A, log2-fold-changes now being normalized to the highest absolute expression value in any time point upon 100 pM stimulation. **D)** 45 immediate early (IEG) target genes. Same as panel A, log2-fold-changes now being normalized to the highest absolute expression value after 45 or 90 min of 100 pM stimulation**. E)** 315 inconsistent target genes. Log2-fold-changes are shown without normalization. **F)** 723 remaining ungrouped genes, whose kinetics are rejected by the kinetic model (Figure 3) and could not be assigned to any of the groups in A-E. Log2-fold-changes are shown without normalization.

## S3

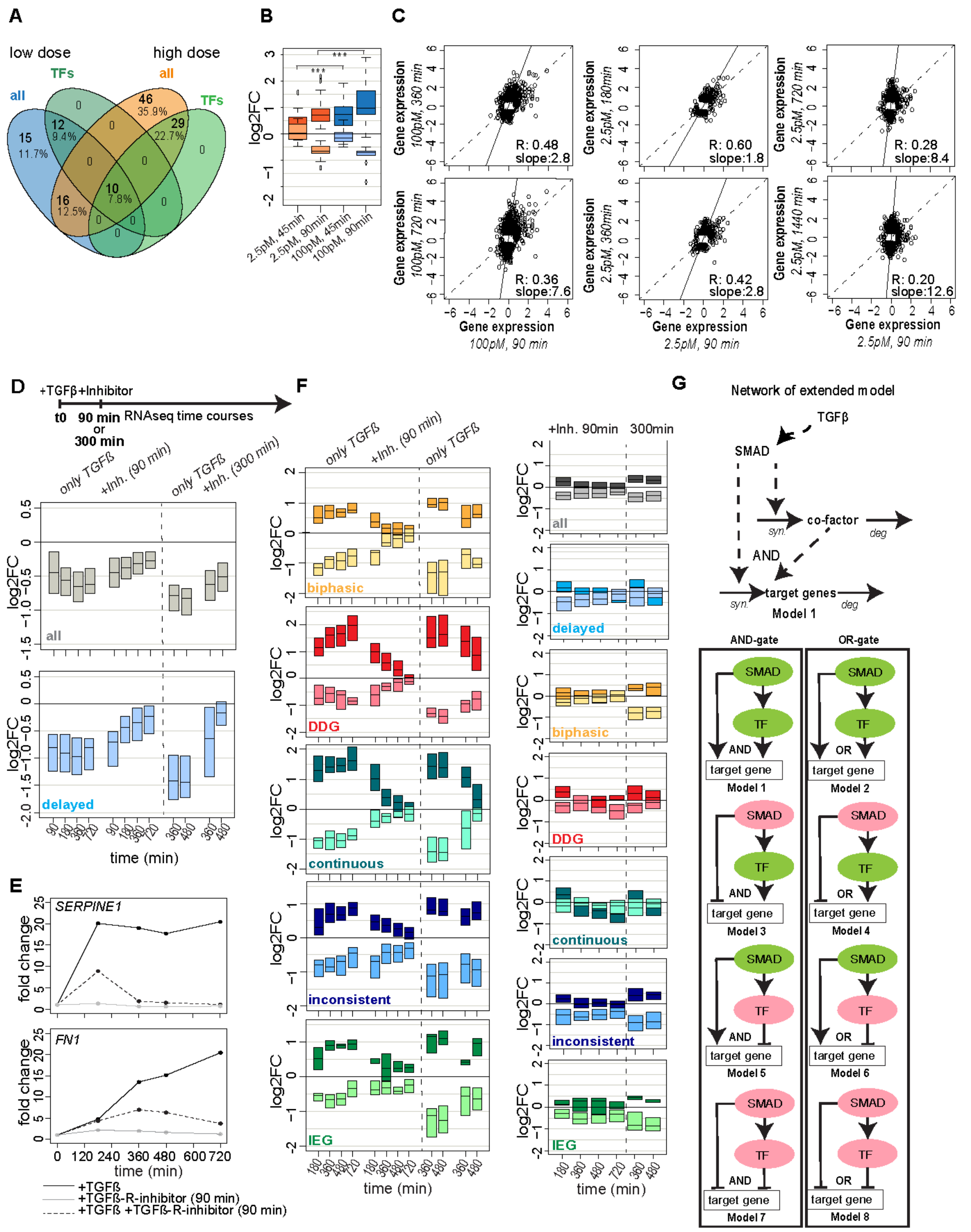


**S3: Evidence for feedforward regulation in TGFβ-dependent target gene expression,** *related to main Figure 4*

**A)** Transcription factors are regulated early after TGFβ stimulation. Venn diagram illustrating the count of all differentially expressed target genes (adj. p-value <0.01, abs. FC > 1.5) and transcription factors (TFs) during the early stages (45/90 min) following low and high dose stimulation. Among the differentially expressed genes, 22 out of 53 are TFs in response to low-dose stimulation (2.5 pM), while 39 out of 105 are TFs in response to high-dose stimulation (100 pM).

**B)** Stronger regulation of TF levels upon high-dose stimulation (100 pM TGFβ) when compared to low-dose stimulation (2.5 pM TGFβ). Boxplot shows log2 fold change of 41 upregulated TFs and 10 down regulated TFs upon 2.5 pM (blue) and 100 pM (orange) TGFβ stimulation. Average expression change of TFs is significantly higher upon high dose (100 pM) stimulation at both time points 45 min and 90 min (45 min: p-value: 0.002, 90 min: p-value: 0.0013).

**C)** Distinct early and late gene expression programs induced by TGFβ. Scatter plots relating log2 fold-changes relative to unstimulated control at various early (x-axis) and late (y-axis) time points as already shown in main Figure 4, F. Each dot represents a significant differentially expressed gene in at least one stimulation condition (100 pM: 90 vs. 360 min: 1112 genes, 90 vs. 720 min: 2930 genes, 2.5 pM: 90 vs. 180min: 389 genes, 90 vs. 360 min: 461 genes, 90 vs. 720 min: 1234 genes, 90 vs. 1440 min: 1496 genes). Lines: bisecting (dashed) and linear fit to the data (solid), with the slope and correlation coefficient indicated on the bottom right.

**D)** RNA sequencing after TGFβ-receptor inhibitor treatment confirms SMAD-dependency of late down regulated target genes. Cells were stimulated with 100 pM TGFβ at t=0 and the TGFβ-receptor inhibitor SB431542 was applied 90 or 300 min afterwards. Genome-wide RNA sequencing was performed at the indicated time points, either in cells treated with TGFβ only (left) or after TGFβ-receptor inhibitor addition (right). The boxes show that the expression distribution (median with lower and upper quartile) of all downregulated (log2FC < - 0.58, adj. p-value < 0.01) TGFβ target genes (top: number of all down regulated: +90 min: 860/ +300 min: 690) or those belonging to the delayed gene expression group (bottom: delayed downregulated: +90 min: 63, + 300 min: 64) increases over time upon inhibitor treatment.

**E)** Time course expression of SERPINE1 and FN1 upon TGFβ stimulation (100 pM, black line), TGFβ stimulation followed by TGFβ-receptor inhibitor treatment 90 min after initial stimulation (100 pM TGFβ, 10 µM SB431542, dashed black line) and TGFβ-receptor inhibitor treatment control (10 µM SB431542, grey line) are shown. **F)** TGFβ-R-inhibitor treatment influences gene groups defined in main Figure 4 (biphasic, dose-discriminating (DDGs), continuous, inconsistent and IEG). The sustained differential expression in biphasic kinetics declines for upregulated target genes: (+ 90 min: 28 genes, +300 min: 22 genes), and increases for biphasic down-regulated target genes (+90 min: 19 genes, +300 min: 18 genes). Late expression boost of DDGs drops for upregulated genes (+90 min: 55 genes, +300 min: 45 genes) and increases for down regulated genes: (+90 min: 17 genes, +300 min: 4 genes). The fast and sustained differential expression of continuous target genes declines in upregulated ones (+90 min: 49 genes, +300 min: 37 genes) and increases in down regulated ones (+90 min: 5 genes, +300 min: 4 genes). The same holds for IEG upregulated genes (+90 min: 5 genes, +300 min: 5 genes) and IEG down regulated genes (+90 min: 8 genes, +300 min: 13 genes) as well as inconsistent genes upregulated: (+90 min: 25 genes, +300 min: 32 genes) and downregulated genes (+90 min: 31 genes, +300 min: 50 genes). Same number of genes shown in Figure D and E, are shown in TGFβ-R-inhibitor treatment control. Inhibitor treatment alone does not result in strong effects.

**G)** Network structure of extended model version. TGFβ activates the SMAD complex that always induces the expression of a transcription factor (TF), which in turn regulates target genes jointly with SMAD, functioning either as a repressor (red) or activator (green) with AND- or OR-gate logic. In Model 1, both SMAD, and the TF function as activators (green) within an AND-gate logic. Model 2 integrates these two activators using an OR-gate logic. In Models 3 and 4 SMAD serves as an inhibitor (red), while the TF acts as an activator (green) in an AND- and OR-gate setting, respectively. In Models 4 and 5 SMAD acts as an activator (green), whereas the TF acts as repressor (red) in an AND- and OR- gate setting. Models 7 and 8 incorporate both SMAD and TF as repressors (red) in an AND-and OR-gate logic, respectively.

**S4**

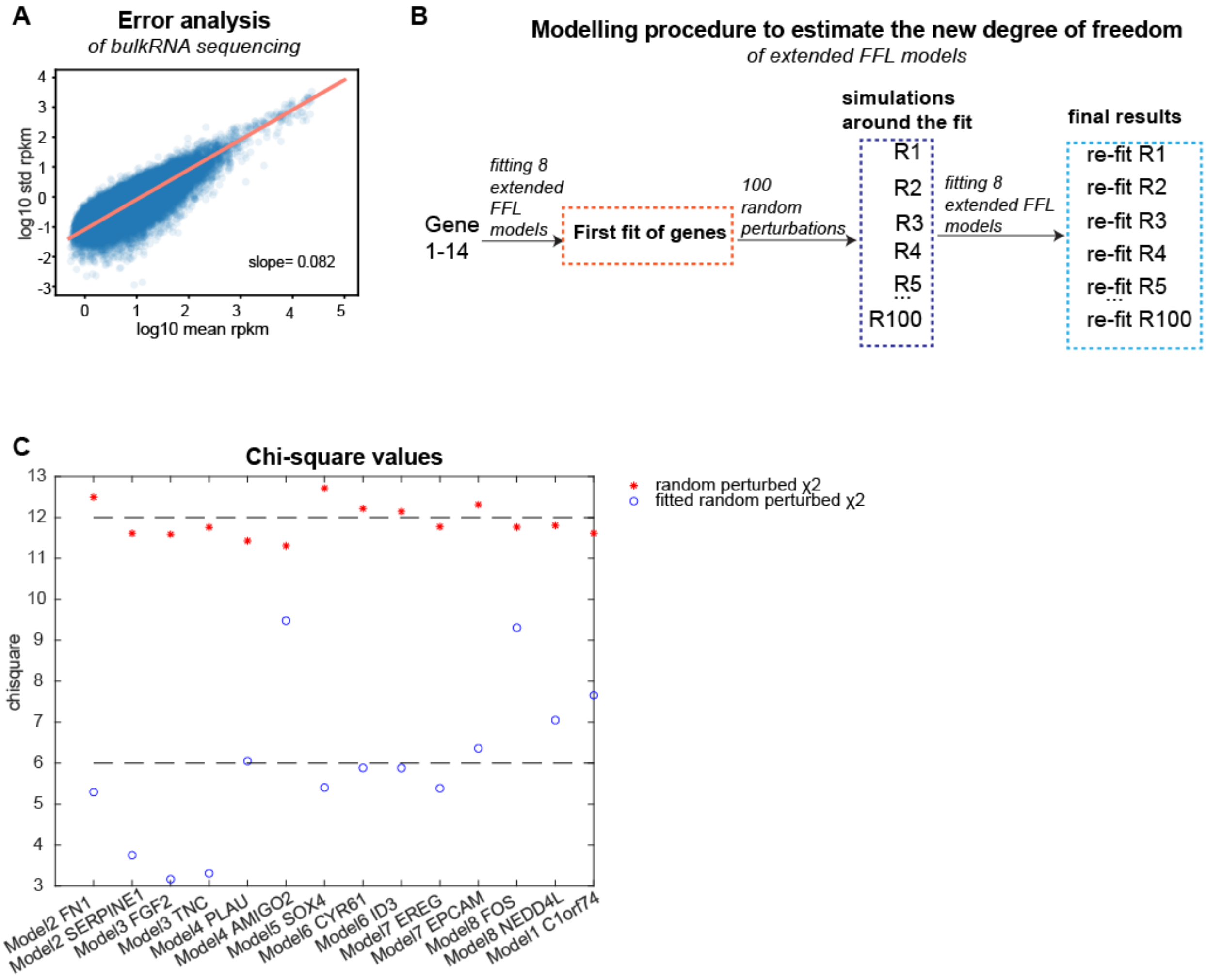


**S4: Model evaluation analysis**, *related to Methods (Model extension by feed-forward loops)*

**A)** Error analysis of time course bulk RNA sequencing data (Figure 1). Mean RPKM values (a) of all 4823 differentially expressed target genes are plotted against the standard deviation (b). Each dot represents a gene. The calculated slope (m) of the scatter plot has a value of 0.082 and is used for relative error calculation (see Methods).

**B)** To determine the effective degrees of freedom of the extended FFL models, we generated synthetic data from best fit and calculated model-intrinsic $\chi^2$ by refitting. 14 genes were selected from all transcription dynamic groups and each of them was fitted to the 8 FFL models. 100 samples were generated by adding normally distributed noise (11% standard deviation) to the best fitted time courses (R1 – R100). The 8 FFL models were refitted to the 100 samples and the mean of the resulting $\chi^2$ values was used to calculate the model-intrinsic degrees of freedom.

**C)** The degree of freedom was calculated from $\chi^2$ values before and after model refit to the synthetic data by $\chi_r{}^2 - \chi_{fr}{}^2 = 12 - 6 = 6$ (see Methods).

**S5**

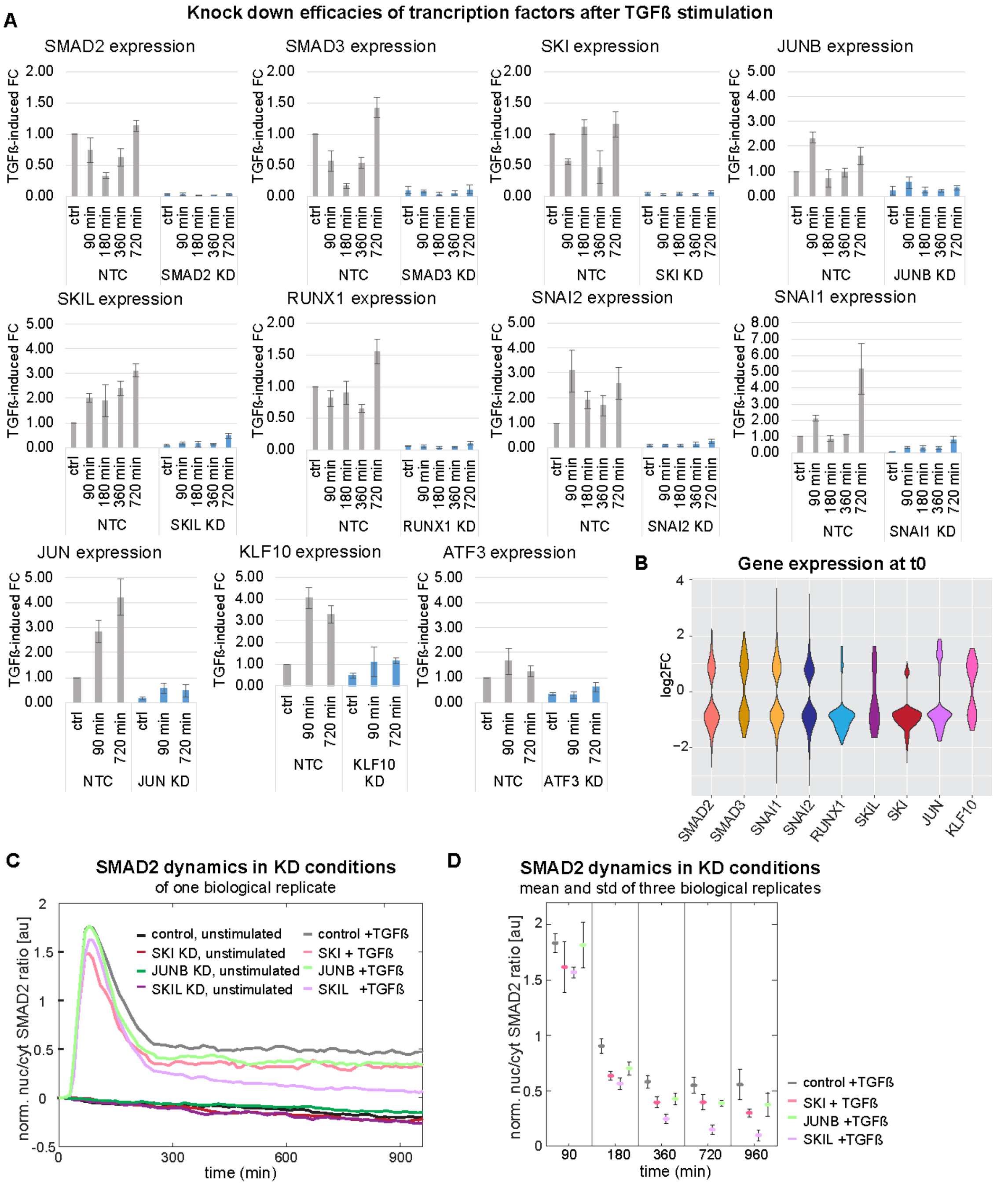


**S5: Knockdown effects on basal gene expression and nuc/ cyt SMAD2 ratio,** *related to main Figure 5*

**A)** Transcription factor knockdown efficiencies at various time points of TGFβ stimulation were evaluated using qPCR. The provided data displays three replicates per condition (control/90/180/360/720 min) for MCF10A cells with knockdowns of SMAD2, SMAD3, JUNB, SKI, SKIL, RUNX1, SNAI1, and SNAI2, along with the SEM. For transcription factors with minimal impact on global gene expression changes JUN, KLF10, and ATF3 only three replicates of control/90/720 min conditions are presented for both control and knockdown MCF10A cells.

**B)** Violin plot shows basal gene expression effects of transcription factor KD. Gene expression of differentially expressed genes (adj. p-value < 0.01, abs. FC > 1.5) without TGFβ stimulation for SMAD2 (485 genes), SMAD3 (182 genes), SNAI1 (846 genes), SNAI2 (1254 genes), RUNX1 (99 genes), SKIL (23 genes), SKI (268 genes), JUN (68 genes), and KLF10 knockdown (86 genes). There are no sign. differentially expressed genes at t0 in JUNB and ATF3 KD.
**C)** Transcription factor KD effects on SMAD signaling are weak. Normalized nuc/cyt SMAD2 ratio upon unstimulated and stimulated (100 pM) JUNB, SKI and SKIL knockdown MCF10A cells of one representative replicate.
**D)** Mean values and standard deviation of norm. nuc/cyt SMAD2 ratio from three biological replicates at specific time points post-stimulation are shown. Plot comprises stimulated (100pM TGFβ) control, SKI KD, JUNB KD and SKIL KD cells. The number of cells used for analysis is shown in Table S7.

**S6**

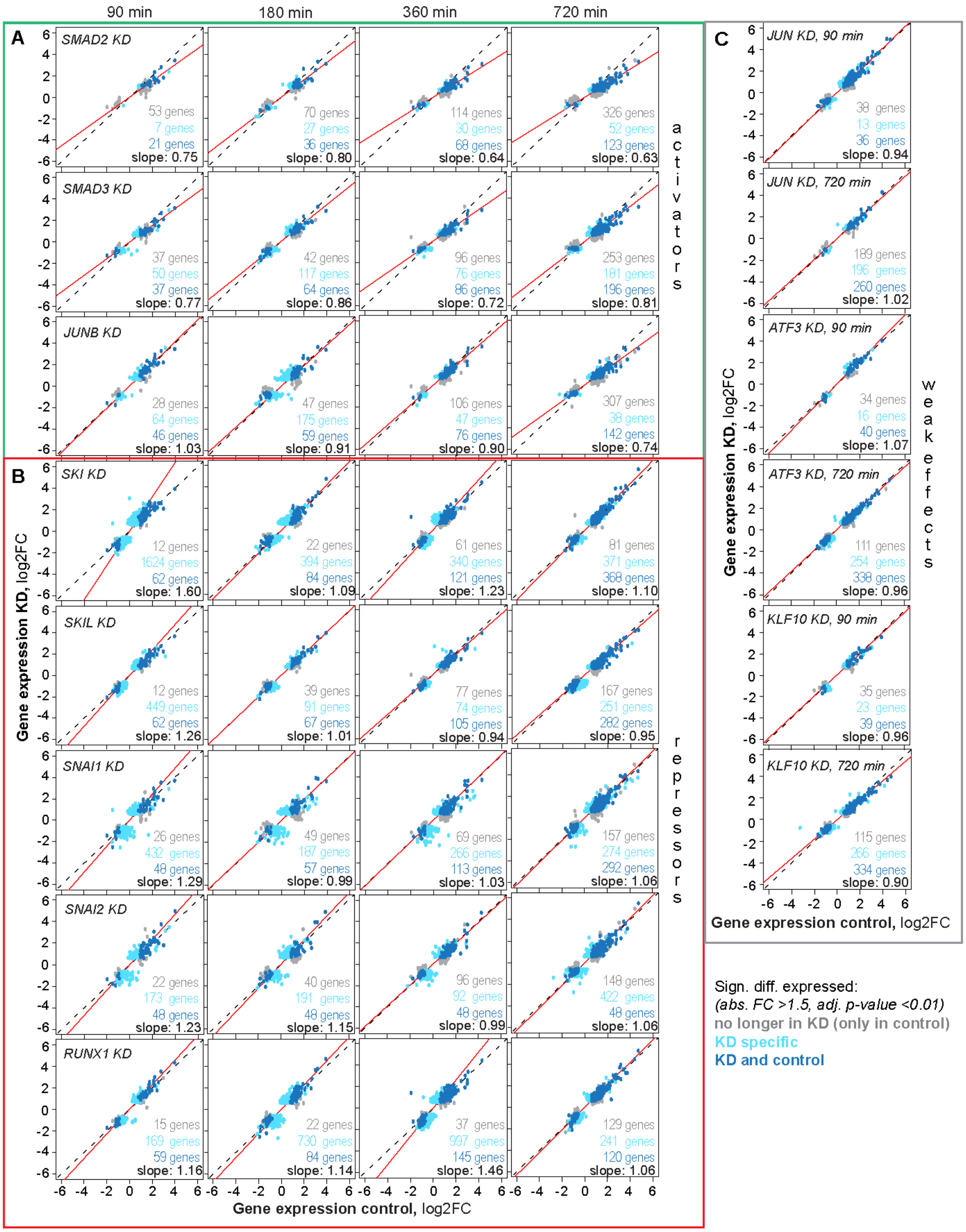


**S6: Activators and repressors of SMAD-mediated gene regulation revealed upon transcription factor KD,** *related to main Figure 5*

**A), B), C)** Scatter plots show TGFβ-induced global gene expression changes in control vs. TF KD 90/ 180/ 360/ 720 min after high dose TGFβ (100 pM) stimulation. The slope of a linear trend line (red line) across all genes determines the strength of A) the activators (top, green rectangle) or B) repressors (bottom, red rectangle). Genes significantly differentially expressed (adj. p-value < 0.01, abs. FC > 1.5) in control only, KD only or control and KD are colored in grey, light turquoise, and dark turquoise, respectively. C) JUN, ATF3 and KLF10 only have weak effects on global gene expression change as highlighted by the given slope.

**S7**

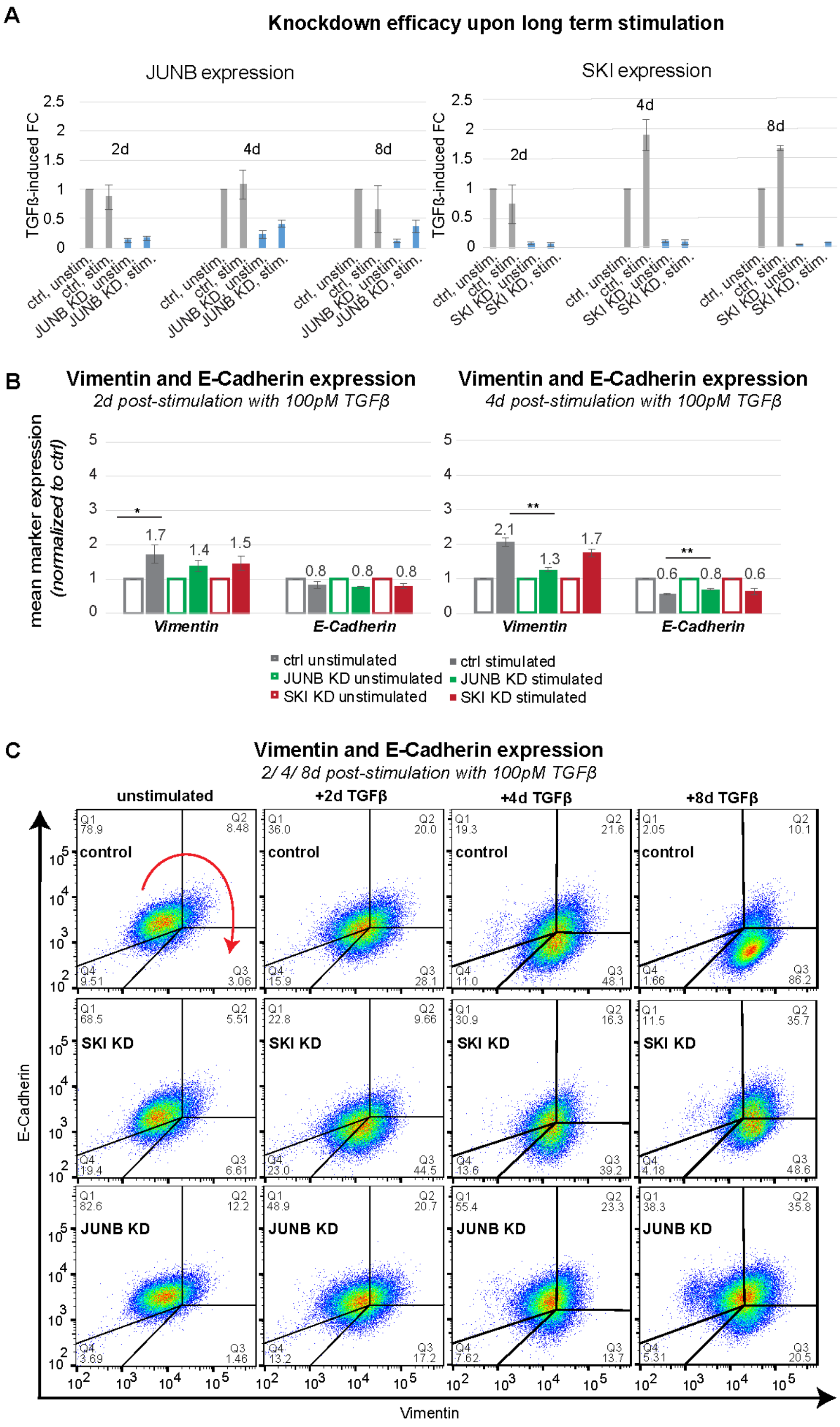


**S7: Phenotypic effects of JUNB and SKI knockdown,** *related to main Figure 5, E*

**A)** Knockdown efficacies of JUNB and SKI KD for 2/ 4 and 8 d normalized to unstimulated control. **B)** The average expression levels of Vimentin and E-Cadherin 2 and 4 d post-stimulation with 100 pM TGFβ were assessed by flow-cytometry. While there is no significant change in marker expression 2 d after stimulation, 4 d post-stimulation Vimentin levels are significantly decreased

and E-Cadherin levels are significantly increased in JUNB KD compared to control. Shown are means of three replicates +/- SEM.
**C)** Density plots of Vimentin and E- Cadherin expression of unstimulated and long-term stimulated (2/4/8 d) control, SKI and JUNB KD MCF10A cells, assessed by flow cytometry, are shown. Red arrow indicates cellular movement in epithelial-mesenchymal-transition (EMT) from Q1 (E-Cadherin$^{high}$, Vimeintin$^{low}$ ) to Q3 (E-Cadherin$^{low}$, Vimentin$^{high}$). Numbers indicate percentage of cells in each quarter (Q1-Q4).

**S8**

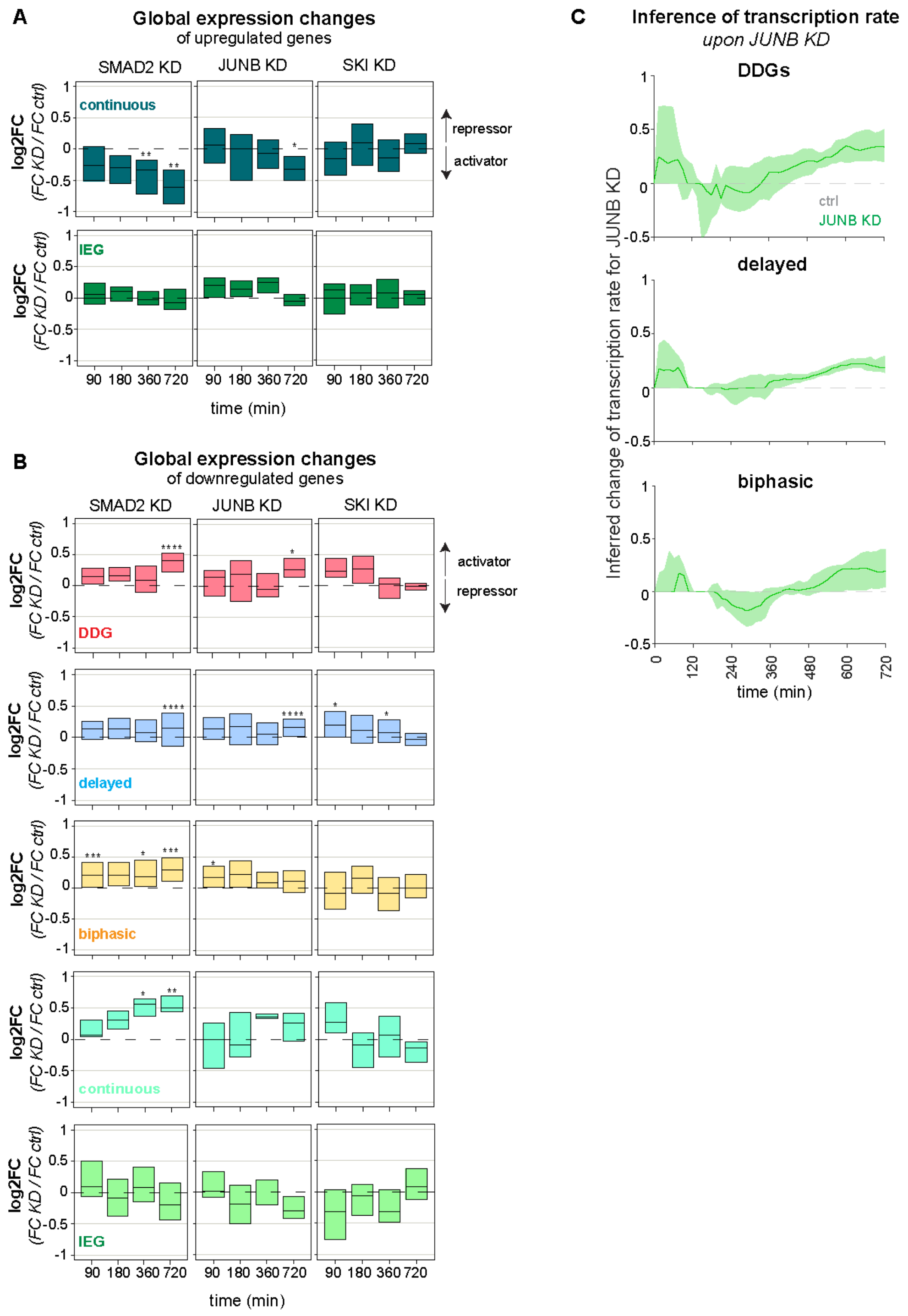


**S8: Global expression changes of dynamic gene groups upon SMAD2, JUNB and SKI KD,** *related to main Figure 6*

**A)** JUNB and SMAD2 KD globally boosts late expression of continuous target genes (turquoise, 56 genes). Immediate early genes remain unchanged from KDs (green, 19 genes). Boxes show changes of TGFβ-induced gene expression upon KD (log2FC = (KD at t=x/ KD at t=0) / (control at t=x/ control at t=0)) as a distribution across all genes belonging to the indicated gene set (median with lower and upper quantile shown).

**B)** For rejected downregulated target genes, SMAD2 KD tends to induce gene expression changes already at early time points as shown for biphasic target genes (yellow, 48 genes) and IEG target genes (green, 25 genes) and late time points for DDGs (red, 12 genes), delayed genes (blue, 182 genes), biphasic and continuous (turquoise, 7 genes). JUNB KD specifically acts at the late 720 min after TGFβ stimulation for the DDGs and delayed target genes and SKI KD tends to change the expression of a subset of genes early after 90/ 180/ 360 min (*p<=0.05, **p<=0.01, ***p<=0.001, ****p<=0.0001).

**C)** Transcription-rate inference on SMAD-downregulated genes. Time-dependent transcription rates ($v(t)$) were inferred from RNA sequencing data for JUNB KD and control cells. The changes in transcription rate ($\Delta v = v_{KD} - v_{control}$) were calculated for downregulated DDGs (top), delayed (middle) and biphasic (bottom) genes. Lines: median of $\Delta v$ in each gene group. Shades: bootstrapped confidence bands (2000 bootstrap samples).

**Table S1: Absolute number of EMT and non-EMT genes in DDGs and non-DDGs,** *related to main Figure 2,* The EMT gene list was curated, and the following target genes related to EMT were added: *WNT9A* (Gasior *et al.*, 2017; Zhang *et al.*, 2018), *AMIGO2* (Kanda *et al.*, 2017; Tanio *et al.*, 2021), *COL4A1* (Miyake *et al.*, 2017; Cui, Shan and Qiao, 2022; Tian *et al.*, 2023), *EPCAM* (Hyun *et al.*, 2016; Sankpal *et al.*, 2017), *FAP* (Wu *et al.*, 2020; Ping *et al.*, 2023), *LAMB3* (Liu *et al.*, 2019; Zhang *et al.*, 2019), *LAMA3* (Huang and Chen, 2021; Islam *et al.*, 2023), *PLAU* (Chen *et al.*, 2021; Wu *et al.*, 2022), *ITGB6* (Thomas, Nyström and Marshall, 2006; Zheng *et al.*, 2021), *SULF2* (Vicente *et al.*, 2015; Tao *et al.*, 2017), *TEAD2* (Diepenbruck *et al.*, 2014), *LTBP2* (Wan *et al.*, 2017; Wang *et al.*, 2018).

| | **EMT genes** | **Non-EMT** |
|---|---|---|
| **DDGs** (86) | 29 | 57 |
| **Non-DDGs** (4737) | 142 | 4595 |
| ***total*** | *171* | *4652* |

**Table S2.** GSEA of DDGs considering KEGG (Kyoto Encyclopedia of Genes and Genomes) and REAC (Reactome) gene sets with adjusted p-value< 0.01, *related to main Figure 2, see supplementary method: GSEA of DDGs*

| **Source and term name** | **Adjusted p-value** | **Number of DDGs intersecting with gene set** | **Gene symbols** |
|---|---|---|---|
| **KEGG**, ECM-receptor interaction | 1.1886E-05 | 7 | *COL4A1, FN1, ITGB6, LAMA3, LAMB3, LAMC2, TNC* |
| **KEGG**, HPV infection | 1.6827E-05 | 11 | *COL4A1, FN1, FZD2, ITGB6, LAMA3, LAMB3, LAMC2, LFNG, PIK3CD, TNC, WNT9A* |
| **KEGG**, Focal adhesion | 0.00027557 | 8 | *COL4A1, FN1, ITGB6, LAMA3, LAMB3, LAMC2, PIK3CD, TNC* |
| **KEGG**, Small cell lung cancer | 0.00030181 | 6 | *COL4A1, FN1, LAMA3, LAMB3, LAMC2, PIK3CD* |
| **KEGG**, Amoebiasis | 0.00051999 | 6 | *COL4A1, FN1, LAMA3, LAMB3, LAMC2, PIK3CD* |
| **KEGG**, PI3K-Akt-Signaling pathway | 0.01610993 | 8 | *COL4A1, FN1, ITGB6, LAMA3, LAMB3, LAMC2, PIK3CD ,TNC* |
| **REAC**, Non-integrin membrane-ECM interactions | 0.00040216 | 6 | *COL4A1, FN1, LAMA3, LAMB3, LAMC2, TNC* |
| **REAC**, Anchoring fibril formation | 0.00056117 | 4 | *COL4A1, LAMA3, LAMB3, LAMC2* |
| **REAC**, ECM proteoglycans | 0.00185292 | 6 | *COL4A1, FN1, ITGB6, LAMA3, SERPINE1, TNC* |
| **REAC**, Extracellular matrix organization | 0.00301222 | 10 | *COL4A1, FN1, ITGB6, LAMA3, LAMB3, LAMC2, LTBP2, LTBP3, SERPINE1, TNC* |
| **REAC**, Signaling by TGF-beta Receptor Complex | 0.00535432 | 6 | *ITGB6, LTBP2, LTBP3, PMEPA1, SERPINE1, SKIL* |
| **REAC**, MET promotes cell motility | 0.00336972 | 4 | *FN1, LAMA3, LAMB3, LAMC2* |

**Table S3.** Number of differentially regulated genes with phenotypic association, *see Methods part: Bioinformatics analysis of bulk RNA sequencing data described or rejected by the simple model, related to main Figure 3*

| | **all genes** | **EMT-related** | **cell cycle-related** |
|---|---|---|---|
| **differentially regulated by TGFβ** | 4823 | 172 | 391 |
| **described by activation model** | 919 (19 %) | 41 (24 %) | 52 (13 %) |
| **described by inhibition model** | 2160 (45 %) | 37 (22 %) | 183 (46 %) |
| **rejected** | 1744 (36 %) | 93 (54 %) | 156 (39 %) |

**Table S4.** Filter settings defining numbers of genes belonging to dynamic gene groups, *related to main Figure 4*
For a gene to belong to a group its RNA sequencing time course had to fulfill the indicated filter settings upon high dose (100 pM) TGFβ stimulation. For instance, to classify a gene with delayed kinetics its absolute (abs.) fold change (FC) needed to be greater than 2.5 in at least one of the time points 180 or 360 or 720 or 1440 min post-stimulation. Simultaneously, the absolute fold change for early time points (45 min and 90 min) needed to be smaller than 1.5. For a gene to belong to the inconsistent gene group its absolute fold change needed to higher than 1.5 and simultaneously lower than 0.8 in at least one of the time points (45/90/180/360/720/1440 min).

| **kinetics** | **Selected filter settings** | **Number of genes rejected *(1744)*** | **Number of genes described by inhibition model *(2160)*** | **Number of genes described by activation model *(919)*** |
|---|---|---|---|---|
| **I) Delayed** | *180 min OR 360 min OR 720 min OR 1440 min abs. FC >2.5, 45 min AND 90 min abs. FC < 1.5* | 483<br>↓189 ↑294 | 277 | 131 |
| **II) Inconsistent** | *45 OR 90 OR 180 OR 360 OR 720 OR 1440 min FC > 1.5*<br><br>*AND*<br><br>*45 OR 90 OR 180 OR 360 OR 720 OR 1440 min FC < 0.8* | 315 | 3 | 6 |
| **III) Biphasic** | *45 OR 90 min abs. FC >1.5,*<br>*abs. FC 180 min < 90 min, 180 min abs. FC < 1.5,*<br>*360 OR 720 OR 1440 min abs. FC >1.5*<br><br>*180 min abs. FC >1.5,*<br>*abs. FC 360 min < 180 min, 360 min abs. FC < 0.58,*<br>*720 OR 1440 min abs. FC > 1.5* | 112<br>↓48 ↑64 | 16 | 22 |
| **IV) Continuous** | *90 min AND 180 min AND 360 min AND 720 min AND 1440 min abs. FC >1.5* | 66<br>↓8 ↑58 | 1 | 26 |
| **V) Immediate early** | *45 min OR 90 min abs. FC >1.5, 180 min AND 360 min OR 720 min AND 1440 min abs. FC < 1.5* | 45<br><br>↓25 ↑20 | 1 | 5 |

**Table S5.** Absolute number and percentages of genes classified in different dynamical gene groups described by the extended model versions model 1-8. There are rejected genes grouped in more than one gene group. In total, 1577 out of 1744 target genes are described by the extended model versions, *related to main Figure 4, D, E, see Methods part: Feed-Forward loops equation, FFL model fitting, FFL model fitting selection*

| | **Model 1** | **Model 2** | **Model 3** | **Model 4** | **Model 5** | **Model 6** | **Model 7** | **Model 8** | **Total (%)** |
|---|---|---|---|---|---|---|---|---|---|
| **Biphasic** | | | | | | | | | |
| (absolute) | 1 | 19 | 20 | 9 | 5 | 5 | 14 | 1 | 74 |
| (%) | 0.9 | 17.0 | 17.9 | 8 | 4.5 | 4.5 | 12.5 | 0.9 | 66.2 |
| **Continuous** | | | | | | | | | |
| (absolute) | 4 | 18 | 4 | 17 | 4 | 3 | 2 | 2 | 54 |
| (%) | 6.1 | 27.3 | 6.1 | 25.8 | 6.1 | 4.5 | 3 | 3 | 81.9 |
| **Delayed** | | | | | | | | | |
| (absolute) | 39 | 42 | 99 | 102 | 2 | 6 | 79 | 66 | 435 |
| (%) | 8.1 | 8.7 | 20.5 | 21.1 | 0.4 | 1.2 | 16.4 | 13.7 | 89.8 |
| **Inconsistent** | | | | | | | | | |
| (absolute) | 0 | 42 | 107 | 102 | 0 | 3 | 17 | 0 | 271 |
| (%) | 0 | 13.3 | 34 | 32.4 | 0 | 1 | 5.4 | 0 | 86.1 |
| **IEG** | | | | | | | | | |
| (absolute) | 0 | 6 | 11 | 5 | 0 | 7 | 1 | 0 | 30 |
| (%) | 0 | 13.3 | 24.4 | 11.1 | 0 | 15.6 | 2.2 | 0 | 65.2 |
| **DDGs** | | | | | | | | | |
| (absolute) | 6 | 4 | 6 | 17 | 1 | 2 | 6 | 4 | 46 |
| (%) | 10.9 | 7.3 | 10.9 | 30.9 | 1.8 | 3.6 | 10.9 | 7.3 | 83.6 |
| **ungrouped** | | | | | | | | | |
| (absolute) | 10 | 72 | 226 | 198 | 14 | 66 | 168 | 53 | 807 |
| (%) | 1.2 | 8.4 | 26.4 | 23.2 | 1.6 | 7.7 | 19.6 | 6.2 | 94.4 |
| **Total** | | | | | | | | | |
| (absolute) | 54 | 183 | 428 | 409 | 25 | 88 | 268 | 122 | **1577** |
| (%) | 3.4 | 11.6 | 27.1 | 25.9 | 1.6 | 5.6 | 17 | 7.7 | **90.4** |

**Table S6.** Quantified slopes and Pearson correlation coefficient (R) of scatter plots (gene expression control vs. gene expression KD) for each time point, *related to main Figure 5, see Methods part: Assessing impact of co-factor KD on target gene expression (slope quantification)*

| Factor KD | Time point post-stimulation (min) | Slope control vs. KD (DESeq2) | Slope control vs. KD (DESeq2) | Variance confidence interval | Pearson Correlation coefficient (R), ctrl vs. KD |
|---|---|---|---|---|---|
| **SMAD2** | 90 | 0.75 | 0.74 | [0.74, 1.32] | 0.94 |
| | 180 | 0.80 | 0.80 | [0.60 1.60] | 0.95 |
| | 360 | 0.64 | 0.64 | [0.97 1.02] | 0.93 |
| | 720 | 0.63 | 0.61 | [0.90, 1.11] | 0.92 |
| **SMAD3** | 90 | 0.77 | 0.74 | [0.87, 1.14] | 0.94 |
| | 180 | 0.86 | 0.82 | [0.60 1.56] | 0.96 |
| | 360 | 0.72 | 0.73 | [0.98 1.02] | 0.94 |
| | 720 | 0.81 | 0.76 | [0.88, 1.13] | 0.92 |
| **SNAI1** | 90 | 1.29 | 1.10 | [0.93, 1.07] | 0.87 |
| | 180 | 0.99 | 0.93 | [0.92 1.08] | 0.88 |
| | 360 | 1.03 | 0.89 | [0.96 1.05] | 0.88 |
| | 720 | 1.06 | 1.03 | [0.88, 1.12] | 0.93 |
| **SNAI2** | 90 | 1.23 | 1.17 | [0.88, 1.13] | 0.88 |
| | 180 | 1.15 | 1.10 | [0.87 1.14] | 0.90 |
| | 360 | 0.99 | 1.01 | [0.92 1.09] | 0.92 |
| | 720 | 1.06 | 1.02 | [0.94, 1.06] | 0.93 |
| **RUNX1** | 90 | 1.16 | 1.13 | [0.94, 1.06] | 0.95 |
| | 180 | 1.14 | 1.00 | [0.91 1.10] | 0.94 |
| | 360 | 1.46 | 1.39 | [0.77 1.28] | 0.87 |
| | 720 | 1.06 | 1.03 | [0.93, 1.07] | 0.96 |
| **SKIL** | 90 | 1.26 | 1.21 | [0.86, 1.16] | 0.95 |
| | 180 | 1.01 | 0.96 | [0.93 1.07] | 0.97 |
| | 360 | 0.94 | 0.94 | [0.99 1.00] | 0.95 |
| | 720 | 0.95 | 0.92 | [0.87, 1.14] | 0.95 |
| **SKI** | 90 | 1.60 | 1.58 | [0.41, 1.96] | 0.92 |
| | 180 | 1.09 | 1.00 | [0.85 1.17] | 0.93 |
| | 360 | 1.23 | 1.28 | [0.71 1.4] | 0.91 |
| | 720 | 1.10 | 1.04 | [0.97, 1.03] | 0.97 |
| **JUNB** | 90 | 1.03 | 1.03 | [0.89, 1.12] | 0.94 |

| | | | | | |
|---|---|---|---|---|---|
| | 180 | 0.91 | 0.84 | [0.84 1.18] | 0.89 |
| | 360 | 0.90 | 0.93 | [0.91 1.09] | 0.94 |
| | 720 | 0.74 | 0.73 | [0.78, 1.26] | 0.93 |
| **JUN** | 90 | 0.94 | 0.94 | [0.75, 1.30] | 0.96 |
| | 720 | 1.02 | 0.97 | [0.91, 1.10] | 0.96 |
| **KLF10** | 90 | 0.96 | 0.95 | [0.81, 1.22] | 0.96 |
| | 720 | 0.90 | 0.88 | [0.78, 1.26] | 0.98 |
| **ATF3** | 90 | 1.07 | 1.06 | [0.79, 1.24] | 0.97 |
| | 720 | 0.96 | 0.95 | [0.79, 1.24] | 0.99 |

**Table S7.** Number of cells analyzed for SMAD2 nuc/cyt ratio in control, non-targeting-control and KD MCF10A cells (SKI, SKIL, and JUNB KD), *related to supplementary Figure S5, C, see supplementary method: SMAD2 live-cell imaging upon TGFβ stimulated transcription factor KD cells*

| **Condition** | **replicate** | **Cell number** |
|---|---|---|
| **WT, unstimulated** | R1 | 792 |
| | R2 | 465 |
| | R3 | 562 |
| **NTC, unstimulated** | R1 | 589 |
| | R2 | 446 |
| | R3 | 679 |
| **SKI, unstimulated** | R1 | 386 |
| | R2 | 152 |
| | R3 | 203 |
| **JUNB, unstimulated** | R1 | 539 |
| | R2 | 162 |
| | R3 | 257 |
| **SKIL, unstimulated** | R1 | 206 |
| | R2 | 383 |
| | R3 | 418 |
| **WT, stimulated** | R1 | 642 |
| | R2 | 373 |
| | R3 | 468 |
| **NTC, stimulated** | R1 | 524 |
| | R2 | 339 |
| | R3 | 428 |
| **SKI, stimulated** | R1 | 227 |
| | R2 | 216 |
| | R3 | 205 |
| **JUNB, stimulated** | R1 | 438 |
| | R2 | 243 |
| | R3 | 282 |
| **SKIL, stimulated** | R1 | 562 |
| | R2 | 304 |
| | R3 | 378 |
